# Problems with Estimating Anthesis Phenology Parameters in *Zea mays*: Consequences for Combining Ecophysiological Models with Genetics

**DOI:** 10.1101/087742

**Authors:** Abhishes Lamsal, Stephen M. Welch, Jeffrey W. White, Kelly R. Thorp, Nora Bello

## Abstract

Ecophysiological crop models encode intra-species behaviors using constant parameters that are presumed to summarize genotypic properties. Accurate estimation of these parameters is crucial because much recent work has sought to link them to genotypes. The original goal of this study was to fit the anthesis date component of the CERES-Maize model to 5266 genetic lines grown at 11 site-years and genetically map the resulting parameter estimates. Although the resulting estimates had high predictive quality, numerous artifacts emerged during estimation. The first arose in situations where the model was unable to express the observed data for many lines, which ended up sharing the same parameter value. In the second (2254 lines), the model reproduced the data but there were often many parameter sets that did so equally well (equifinality). These artifacts made genetic mapping impossible, thus, revealing cautionary insights regarding a major current paradigm for linking process based models to genetics.

## Introduction

In the opening sentences of the 1968 book, The Population Bomb, Paul Ehrlich (and his wife Anne, uncredited at publisher behest) wrote, “The battle to feed all of humanity is over. In the 1970s hundreds of millions of people will starve to death in spite of any crash programs embarked upon now” and, in a subsequent chapter, “I don’t see how India could possibly feed two hundred million more people by 1980." Fortunately, research started in Mexico, India and elsewhere by Norman Borlaug before 1968 created high yielding dwarf wheat varieties that, worldwide, are credited with averting one billion deaths from famine. India also introduced IR8, the so-called “miracle rice” developed at the International Rice Research Institute in the Philippines and the predicted human catastrophe was averted.

Nearly 50 years later, the specter of global disruption is again upon us. The challenges today are not only increasing human population (which has doubled since 1970) but emerging concerns like climate change and declining water resources. The confluence of these manifold trends makes finding ways to feed nine billion people by 2050 one of the most pressing issues of our time [1]. However, the annual percentage increase rates for crop yields are only half those required to meet that goal [2].

Beginning over 20 years ago, a paradigm has emerged offering the promise of dramatically accelerating breeding programs via improved phenotype prediction of prospective crop genotypes in novel, time-varying environments subject to sophisticated management practices [3-7]. The basic notion has two parts. The first is to exploit ecophysiological crop models (ECM’s) to describe the intricate, dynamic, and environmentally responsive biological mechanisms that determine crop growth and development on daily or even hourly time scales. The aim is to use highly detailed, nonlinear simulation models to predict the phenotypes of interest within a subsample of possible environments and in-field management options. ECMs, whose origin is often credited to Wit. (1965), encode intra-species behavioral differences in terms of parameters that are intended to summarize genotypic properties. On the strength of that presumption, the constants are termed *genotype-specfc parameters* (GSP’s).

The second part of the paradigm is to use quantitative genetic methods such as genomic prediction [9] to relate the GSP’s to genotypic markers [3]. Next, the outcomes of crosses are estimated by (1) calculating the GSP values that would arise from possible offspring genotypes. These values are then (2) used in ecophysiological model runs to predict the phenotypes in the target population of environments (for which detailed descriptive data must be available). In simplified instances, this approach has seen remarkable success (e.g., [10].

Composed of large coupled sets of continuous-time differential equations, ecophysiological models simulate many interacting processes [11,12] operating in the soil-plant-atmosphere continuum. These processes include physiology (e.g., photosynthesis, respiration, resource partitioning to various plant parts, and growth), phenology (leaf emergent timing, the date of vegetative-to-reproductive development, etc.), as well as chemistry and physics (soil water flows, chemical transformations, energy fluxes, gas exchange, etc.). During simulation runs, model formulas compute instantaneous process rates based on plant status and environmental conditions at each time point. These rates are integrated (*Sensu* calculus) to output time series of dozens of plant variables. The models typically have 10 to 20 GSP’s whose estimates are read from input files at the start of model execution. Numerous other inputs (e.g. soil water holding capacities by layer; measured daily solar radiation, rainfall, maximum and minimum temperatures; etc.) further quantify the physical environment.

The lynchpin of the two-step paradigm is the accurate estimation of the GSP’s so that these can be related to allelic states of the individual lines. Unfortunately, the direct measurement of GSP’s is so time- and resource-demanding as to be infeasible for large numbers of lines. Indirect GSP estimation via model inversion is also challenging because easily-measured plant phenotypes exhibit strong interactions with the environment [13] thus increasing data requirements by necessitating trait measurement in multiple settings [14]. Even so, ecophysiological crop models enjoy extensive global use in areas ranging from global climate change, policy analysis, crop management, etc. Indeed, a Google search on the abbreviations of just two major model systems [namely “DSSAT” [15] and “APSIM” [16]] returned 134,000 hits. Not surprisingly, there is an extensive literature (reviewed briefly below) on ecophysiological model parameter estimation.

Initially, the authors’ intent was to apply the two-step method to anthesis date using data from over 5000 lines comprising the maize nested association mapping population (NAM) [17], which was developed specifically to enable high-resolution studies of trait genetic architectures. Not only is anthesis date a phenotype of major biological significance, but it was also studied in this same panel using conventional statistical genetic methods [18,19]. Our hypothesis was that applying the proposed 2-step paradigm would demonstrate its merit in the specific context of the large data sets increasingly used in crop breeding programs to interrelate genotypes and phenotypes. Contrasting the results of the standard and ecophysiological approaches was expected to be interesting and informative. Granted, the model fitting methods to be used were not novel, but we expected that a further demonstration of their value with data sets much larger than ever used before would have utility.

However, something quite different happened. We discovered modeling issues and estimation artifacts that are of sufficient severity and generality that, if not addressed, are likely to imperil the breeding acceleration paradigm. Therefore, the objectives of this paper were 1) to describe these problems and the methods that revealed them (which can be applied as detection tools in studies of other traits) and 2) to discuss research directions that might ameliorate the problems.

## Background

Numerous optimization methods have been used to estimate parameters for ECM’s. Surprisingly, perhaps the most common approach has been that of trial and error [20], wherein different parameters values are manually tested until an acceptable match between simulated and observed data is found. This approach, of course, becomes highly inefficient as the number of model parameter increases. Thus, numerous off-the-shelf, automated optimization techniques have been developed. Examples include the simplex method [21], simulated annealing [22,23], sequential search software (GENCALC) [24], Uniform Covering by Probabilistic Region (UCPR) [25], particle swarm optimization (PSO) [26], and generalized likelihood uncertainty estimation (GLUE) [27]. While these traditional optimization techniques have advantages, they can be inefficient in terms of runtime and are highly dependent on optimization settings when thousands of combinations of line × planting site-years are involved – a situation that is becoming common in the era of massive genetic mapping populations. The fundamental issue is that, as the number of lines and environments increases, estimating GSP’s for each line independently usually involves highly redundant simulation. To this end, we adapted an algorithm pioneered by Welch et al. (2000) and Irmak et al. (2000), as described in methods section. The approach exhibits particular efficiencies when individual plantings incorporate large numbers of lines and, serendipitously, supports a close examination of the estimation process, itself.

The vast majority of prior ECM parameter estimation studies have been conducted in non-genetic contexts. Against these backgrounds, the sole merit criterion has been the predictive skill demonstrated by the GSP estimates obtained. However, the current setting, however, is markedly different. GSP’s are not just inputs to ecophysiological crop models; GSP’s simultaneously function as the outputs (i.e. dependent) variables of genetic prediction models. As such, GSP’s are at least as closely related to tangible biochemical processes at the molecular level as they are summative of physiological properties (e.g. maximum photosynthetic rates) in higher organizational realms. Therefore, a deeper inspection of their estimation is warranted and two concepts are helpful in achieving the enhanced discernment now required.

We employ the term “expressivity” (and the adjective “expressive”) to describe a model’s innate ability to reproduce a set of observations independent of particular parameter values. An expressive model may fail to replicate data because an unskilled optimizer cannot find a meritorious combination of parameter values. In contrast, a model with low expressivity will fail to fully mimic actual data irrespective of what (biologically or physically reasonable) values are assigned to its parameters. In cases where the latter behavior is detected, remedies will be vigorously sought. However, as shown below, however, systematic gaps in expressivity can coexist even within an overall framework of predictively skilled model performance.

Another model property that has received little attention in previous estimation studies is equifinality. Equifinality describes a situation in which multiple sets of parameter values generate identical model predictions. In statistics, a synonym for “equifinality” is “parameter non-identifiability” [30][31](31). When the only concern is prediction quality and that seems “good enough”, it is easy to consider equifinality a non-problem. However, when parameters are intermediaries rather than just inputs and equifinality exists, it begs the question as to what relationship, if any, putative GSP estimates might bear to allelic states across the genotype? A moment’s reflection shows that equifinality and expressivity are different model properties. The former relates to how many different estimates yield identical predictions; the latter refers to the possible existence of systematic failures of those predictions to mimic observed data.

In this paper, we explore these issues in modeling and estimation using the anthesis phenology component of the CERES-Maize ECM [32-34] and observed dates from multiple plantings of three maize genetics panels totaling nearly 5300 lines. Anthesis initiates the period of grain development and is therefore a critical milestone toward grain yield. As such, it mediates the adaptation of the crop to its environment by determining the relative length of the vegetative and reproductive growth phases and is a key target of breeding programs [18]. (Although at the apical meristem, floral initiation precedes the visible morphological change of anthesis, the linkage between the two is tight enough that we follow common modeling practice and consider them as effectively synonymous.) The genetics of flowering time has been intensively studied in the model plant *Arabidopsis thaliana* where well over 100 influential genes are now known [35]. Indeed, gene expression models of flowering time of *A. thaliana* based on differential equations have been developed [36], and genetically-informed approaches have established the relationships between network-level function and common ecophysiological time formulations [37]. In maize, our understanding of the genetic control on flowering time is more limited but has been advancing in recent years. More than 30 genes have been described and conservation of key features from *A. thaliana* seems apparent (Table 1 in [38]). A quantitative gene network model based on a number of these loci has been published [38].

The general desire within applied quantitative genetics to probe genetic architectures has led to the construction of ever-larger and/or special purpose mapping populations [18]. The maize NAM panel [17] was constructed by making bi-parental crosses between one common parent, B73, and each of a set of 25 other inbreds that collectively encompassed a wide range of maize diversity. Approximately 200 offspring from each of these 25 crosses were then inbred for a number of generations to ensure, to the greatest degree feasible, that the influence of each locus on any trait of interest reflected the contribution of one parent only. Individual plant genotypes produced in this fashion are called “recombinant inbred lines” (RIL’s). Buckler et al. (2009) reported a seminal study of maize anthesis dates using this NAM panel. Demonstrating the power of these lines to finely dissect genetic contributions to traits of interest, they identified 36-39 QTL, where the exact number depended on the analysis method used. Most of loci had small effects but collectively, they explained 89% of total variation in anthesis date.

For the reasons outlined above, accurate prediction of anthesis date is a major target for ecophysiological crop models [25]. However, few studies exist have used large data sets for ECM calibration. Mavromatis et al. (2002) reported 5,109 site-year-line-parameter combinations and Welch et al. (2002) estimated 4,620 site-year-line-parameters. The effort presented herein encompassed 197,964 site-year-line-parameter combinations - to our knowledge, the largest such study ever reported. As the following sections document, it was the sheer scale of this data set and the resulting scatterplots depicting thousands of lines that revealed worrisome issues of equifinality and expressivity that might be overlooked in studies of smaller scale.

## Materials and Methods

### Experimental data

Observations collected on anthesis date for a total of 5266 maize lines were obtained from the Panzea data repository (http://www.panzea.org). The lines used were members of three genetic panels. In particular, 4785 lines were from the 25 RIL panels comprising the maize NAM set described above. Also included were an additional 200 RIL lines commonly referred to as the IBM panel because they originated by 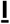 ntermating 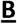73 × **M**o17 [40]. Finally, a maize diversity panel [41] contributed data on 281 additional lines. Various combinations of these lines were grown at six US sites: New York (NY), North Carolina (NC), Illinois (IL), Missouri (MO), Florida (FL) and Puerto Rico (PR), during 2006 and 2007 for a total of eleven site-years. In what follows “NY6” denotes the 2006 planting in New York, respectively by state abbreviation and year for other site-years. Table 1 gives the exact locations of the experimental sites, and the respective sowing dates. The “Total Lines” row of the table gives the number of lines from the three panels that were present in each study. The “Lines with data” row lists the number of lines with available observations on anthesis date. Data on daily maximum and minimum temperatures for each site were provided by the maize NAM collaborators (H. Hung, personal communication, 2010) and did not included metadata on position of the weather stations to the field plots, types and calibration of sensors or types of radiation shields used.

**Table 1.** Sowing dates, geographical coordinates, total number of lines planted and number of lines for which anthesis dates were observed for all site-year combinations used in this study.

|  | NY6 | NY7 | NC6 | NC7 | MO6 | MO7 | IL6 | IL7 | FL6 | FL7 | PR6 |
| --- | --- | --- | --- | --- | --- | --- | --- | --- | --- | --- | --- |
| Sowing Date<br>(DOY) | 128 | 135 | 122 | 120 | 137 | 138 | 128 | 137 | 265 | 280 | 314 |
| Latitude (deg) | 42.73 | 42.73 | 35.67 | 35.67 | 38.89 | 38.89 | 40.08 | 40.08 | 25.51 | 25.51 | 18.00 |
| Longitude (deg) | -76.66 | -76.66 | -78.49 | -78.49 | -92.23 | -92.23 | -88.2 | -88.2 | -80.49 | -80.49 | -66.51 |
| Number of total<br>lines sown | 5478 | 5478 | 5478 | 5478 | 5478 | 5478 | 5478 | 5478 | 5026 | 3753 | 5131 |
| Number of lines<br>with data | 4743 | 5236 | 5236 | 5160 | 3261 | 2555 | 5036 | 5178 | 4943 | 3742 | 4401 |

### CERES-Maize model

The Crop Estimation through Resource and Environment Synthesis (CERES)-Maize model is one of the oldest, most widely used ecophysiological crop models for maize [42]. We used the CERES-Maize version incorporated in CSM (Cropping System Model) 4.5 ([11,15]. The CERES-Maize simulation of development toward anthesis is controlled by a set of GSP’s and environmental inputs [33,34]. Specifically, the GSP’s studied herein were thermal time from emergence to juvenile phase (P1), critical photoperiod (P2O), sensitivity to photoperiods longer than P2O (P2), and the phyllochron interval (PHINT) as measured in thermal time. The duration of Stage 1, the interval from emergence through the end of the juvenile phase, is calculated by accumulating daily thermal time until P1 is reached. Stage 2 follows immediately and lasts until tassel initiation. Stage 2 lasts a minimum of four days when the photoperiod (including civil twilight) is less than P2O. P2 specifies the number of extra days required for every hour by which the photoperiod exceeds P2O. The model continues to accumulate thermal time through Stage 2. The model assumes that (1) there are five embryonic leaves; (2) two new leaves initiate during each phyllochron interval; and (3) that anthesis date, which terminates Stage 3, occurs when all leaves present at the end of Stage 2 (i.e., total leaf number, TOLN) are fully expanded. The date on which this happens is when the ongoing thermal time accumulation reaches TOLN × PHINT.

Thermal time is calculated from inputs of daily maximum and minimum temperatures. Sowing dates (Table 1) determined the time series of weather data that control simulated plant growth and development. The model calculated daily photoperiods from geographic position. Other required model inputs did not affect predicted anthesis dates and were not considered here. For example, the soil water and nutrient balance components of the model do not affect simulated anthesis date in the CERES-Maize model and therefore were not used in this study. The model also requires row spacing and planting depth, which were set to 0.5 m and 2.5 cm, respectively. No tillage, pest, or disease effects were simulated.

### Parameter estimation

#### Search strategy

In the conventional approach to parameter estimation (Fig 1a), an optimizer iterates through a series of trial solutions for which model predictions are generated in each environment. The entire process is repeated for each line. This approach becomes inefficient when many lines are planted together in large experiments and are therefore exposed to identical environments. This is because estimates approaching optimal goodness-of-fit will only emerge in the latter stages of an iterative optimization run. Therefore, the majority of early iterations for each line entail the repeated evaluation of estimates with mediocre predictive ability in the same environment.

**Fig 1.**
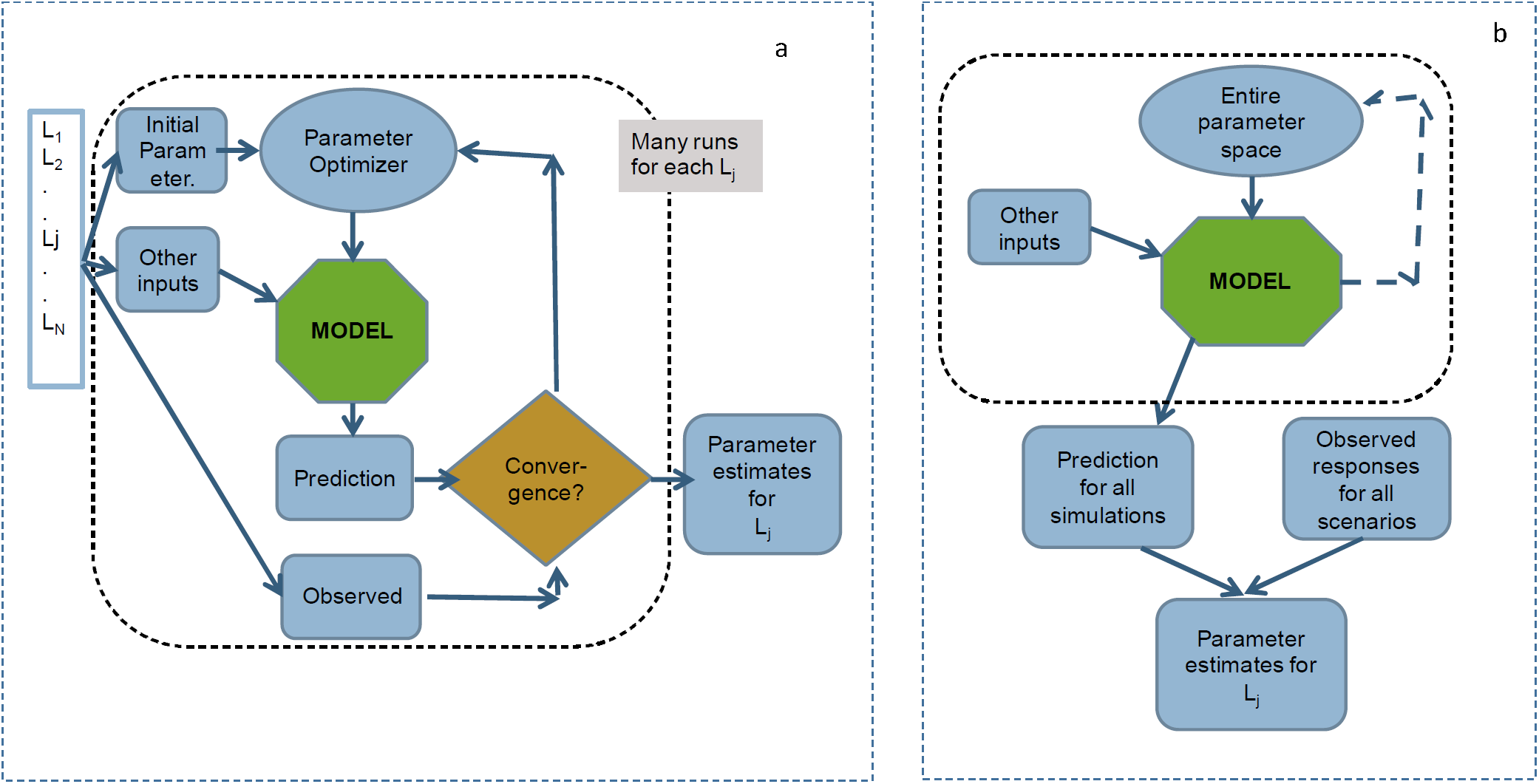
Parameter search strategies. (a) Conventional method (b) Database method. L_l….N_ is the number of lines.

To overcome this problem, we adapted an approach described by Irmak et al. (2000) and Welch et al. (2002, 2000). In their scheme (Fig 1b), model simulations were conducted for each planting across a multidimensional grid of parameter value combinations. The resulting predictions were stored in a database. As a second step, for each line the root mean square error objective function (RMSE; [43] between observed and predicted anthesis day of year was evaluated with respect to all combinations of parameter values across all site-years. That is, for line l,

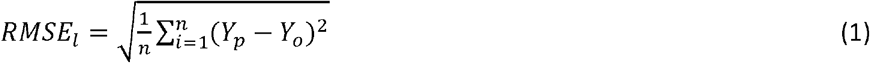

where, *n* is the number of observations for that line (consisting of one observation per site-year combination), and *Y*_p_ (*Y*_o_) is the predicted (observed) anthesis date. The optimizer goal was to minimize the RMSE for each line. If a unique minimum existed, it defined the combination of GSP values that best fit each line. Total computational time was reduced because time-consuming model simulations for each combination of GSP parameter values were only performed once, but those outputs were reused many times in the much faster RMSE calculations. Another benefit is that a combination of GSP values that yielded poor predictability for one variety might perform better for a different line. Additionally, this process ensured that identical parameter combinations were tested for each line, which can aid in comparing the results achieved. Finally, simply by retabulating the database, any number of different optimizations could be performed using different observations, alternative subsets of site-years plantings or combinations of parameter values. The use of alternative objective functions is also possible without requiring additional simulations. Because of the central role played by the database of simulation outputs, we will refer to this scheme as the *database method*.

#### Sampling the model parameter space with sobol sequences

Unlike Irmak et al. (2000) and Welch et al. (2002, 2000) who sampled the parameter space with a rectilinear grid, we employed Sobol sequences so as to avoid the combinatorial explosion in computational requirements that accompany increasing dimensionality. Sobol sequences belong to a family of quasi-random processes designed to generate samples of multiple parameters dispersed as uniformly as possible over the multi-dimensional parameter space [44]. Sobol sequences are specifically designed to generate samples with low discrepancy – that is, a minimal deviation from equal spacing. Unlike random numbers, quasi-random algorithms can effectively identify the position of previously sampled points and fill the gaps between them [45], thus avoiding the formation of clusters. Further, Sobol sequences offer reduced spatial variation compared to other sampling methods (e.g., random, stratified, Latin hypercube; see Fig 2a vs. 2b), make this method more robust [46]. We used a Python-based algorithm to generate a Sobol sequence of quasi-random numbers for calculating 32,400,070 sets of the four CERES-Maize GSP’s, leading to a uniformly-sampled four-dimensional parameter space for P1, P2, P2O, and PHINT. To construct the database, CERES-Maize calculated anthesis date for each GSP combination in each of the 11 site-years - a total of 356,400,770 model runs. Table 2 describes the upper and lower bounds and the number of distinct values obtained for each parameter.

**Fig 2.**
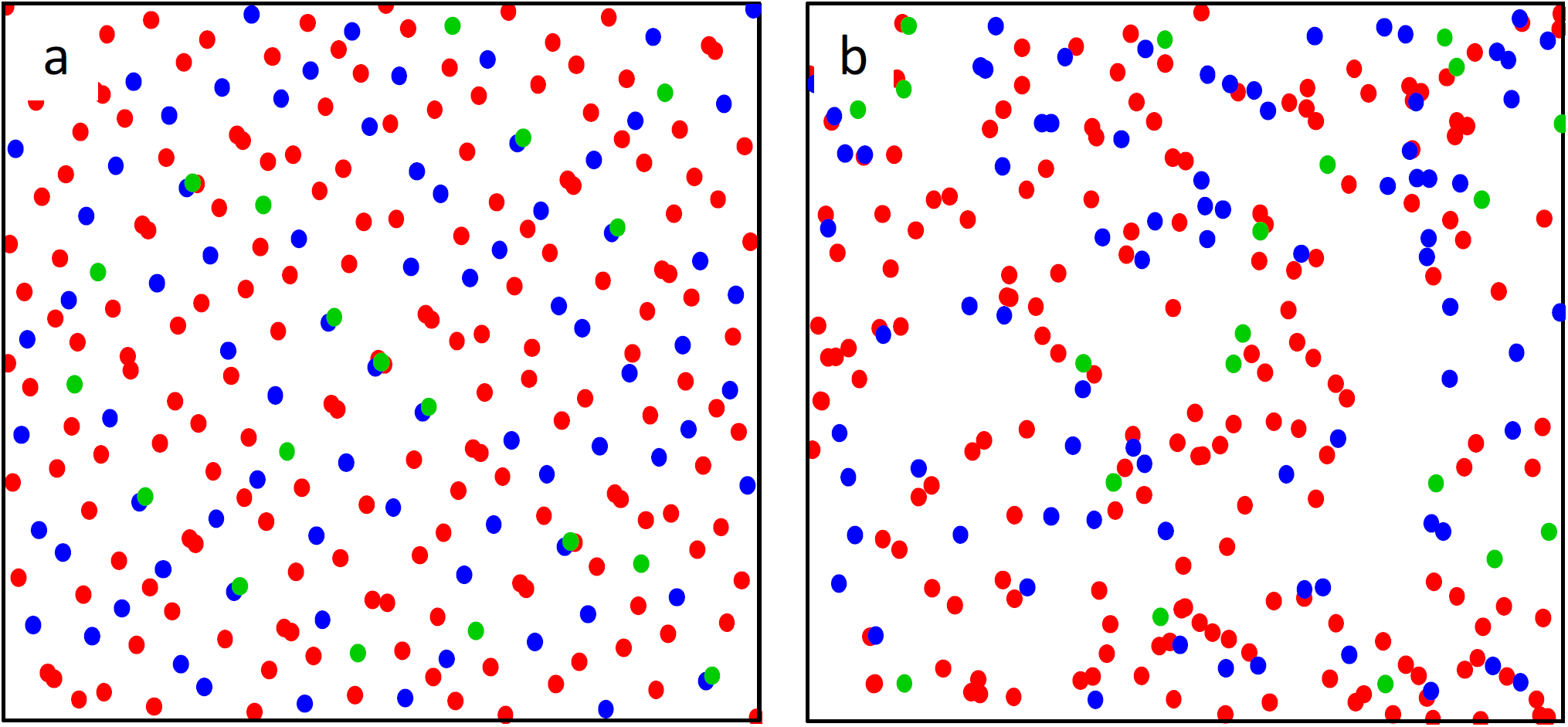
Quasi-random points from two-dimensional Sobol sequence. (a) The first 275 quasi-random points from a two-dimensional Sobol sequence. (b) The first 275 points produced by the commonly used Mersenne twister pseudo-random number generator [47]. The Sobol sequence covers the space more evenly. The first 20 points are green, the next 80 are blue, and the final 175 are red, thus demonstrating Sobol gap filling.

**Table 2.**
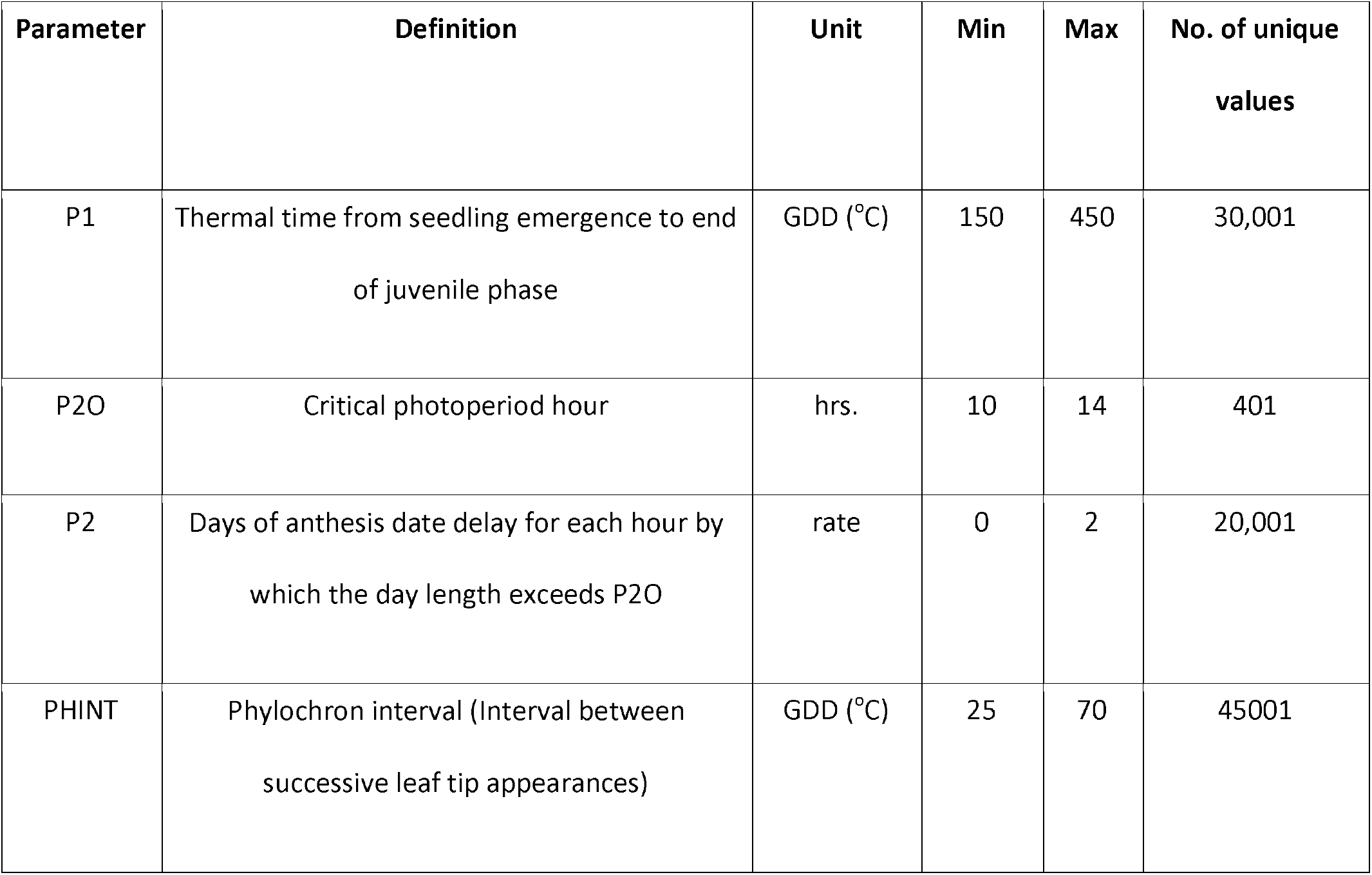
Parameter ranges used in generating sobol sequence.

### High performance computing

The number of model runs was too large for lab-scale computing facilities, so we used the “Stampede” supercomputer at the Texas Advanced Computing Center (TACC) [46]. *In toto*, the CERES-Maize runs required 63,372 CPU-hours, which equates to ca. 176 simulations per second distributed across 112 processors. The predicted anthesis dates were collated and transferred to the “BeoCat” computing cluster at Kansas State University (https://support.beocat.ksu.edu/BeocatDocs/index.php/Compute_Nodes). There, RMSE values were tabulated for each line × parameter value combination across all site-years in which anthesis date was observed. As combinations of GSP values were found that had progressively lower RMSE values, they were recorded by the computer. This process required ca. 15 minutes of wall clock time per line so the total estimation process was completed in ca. 7 h on 200 Xeon E5-2690 cores.

### Assessing estimate properties

#### Equifinality

Equifinality occurs when multiple combinations of parameter estimates generate the same minimal RMSE value, often because they generate identical model predictions [30], in this case identical integer DOY values for anthesis dates. We quantified “equifinality” by defining “number of ties” as the number of Sobol sets of parameter combinations that produced the same optimal RMSE values, minus one. No equifinality is present in a line if there is only one combination of parameter values that minimizes the RMSE. That is, there are zero ties among its estimates. To illustrate the magnitude of the problem and our motivation to study it more closely, we note that 2254 (43%) of the 5266 lines available in the data exhibited equifinality. The worst case was represented by a line that had 1,043,933 distinct combinations of GSP values that produced identical anthesis date predictions, and thus the same RMSE, thereby yielding 1,043,932 ties.

During the database tabulation phase, the values of the “best combination of parameter estimates seen so far” was updated only if its RMSE value was strictly better than all previously evaluated ones. So, when equifinality was present, the final GSP estimate was the first combination of parameter values encountered that had a minimal RMSE value. As a result, some of the analyses described below are sensitive to equifinality, illustrating the fact that subtle optimizer algorithm idiosyncrasies can have marked impacts on the overall results. Such cases are noted explicitly along with the procedures used to mitigate the effects.

### Interrelationships between parameter estimates

Correlations and other relations among parameter estimates are highly important to breeding programs and related simulation studies. When correlations between parameter estimates are present, opportunities exist to select on one plant trait by selecting on a related phenotype instead. Additionally, there have been a number of *in silica* studies where CERES models were used to design crop ideotypes [48,49]. Such efforts find combinations of model parameter values that predict phenotypes well suited to the target population of environments. Once identified, lines with those values become breeding targets. However, a potential pitfall arises if realizing the desired genotype involves changing parameter values in directions contrary to the correlations that exist between them.

For this reason, we explored the pairwise correlation structure of the GSP parameter estimates and generated pairwise scatter plots of their line-specific values. However, the latter revealed a bizarre pattern, the diagnosis of which ultimately led us to the second problem alluded to in the introduction - the inability of the model to reproduce certain observational combinations - and to the methods presented next.

### Model expressivity

A common graphical method to assess the quality of model fit is to plot the predicted vs. observed values (e.g., Fig 3). Such scatterplots can be informative in detecting areas of mismatch between observed and predicted values, thus providing specific characterization of the model’s lack of fit. By definition, each point in the scatterplot corresponds to a prediction that a model is able to make given an optimized set of parameter values. However, an entirely different question is whether there are observations that a given model cannot reproduce using *any* reasonable combination of parameter values? That is, one might seek to assess whether a given model has the requisite expressivity to reproduce the data.

The database approach allows such a question to be addressed using what we term *phenotype* space scatter plots. In such plots, each axis corresponds to a different site-year. The coordinates along the axes represent the observed or predicted anthesis dates for each site-year. Model expressivity is then assessed by comparing the scatter of predicted anthesis date generated from a wide range of GSP value combinations to the scatter of observed values in large data sets. Because equifinality does not affect predictions, this method of evaluating model expressivity is independent of the order in which an optimizer locates points that minimize RMSE values (see the second paragraph in section “Equifinality”).

### Testing for parameter stability across environments

In order for the two-step paradigm outlined in the Introduction to work, the estimates of GSP’s should not vary across the set of environments used to estimate them, a property called “stability” [4]. If GSP estimates did vary across environments, there would be no way to tell what GSP values to input to the ecophysiological model to predict traits whenever daily weather time series or soils differed from those used in the paradigm’s first step. This might seem an insuperable barrier to readers for whom G×E interactions are virtually ubiquitous among quantitative plant phenotypes, but it is not. This is because the raison d’etre of models like CERES-Maize is to explain crop variety × environment interactions mechanistically based on physiological principles.

Many GSP’s, including the ones in this study, explicitly relate plant behaviors (e.g., development toward anthesis) to environmental variables (e.g., temperature and photoperiod in the current case). Modelers assert that GSP’s are properties of the individual lines (i.e., stable) and, therefore, by implication, have a genetic basis because genotypes do not change with the environment. Over time, it is thus expected that research will mechanistically link at least some GSP’s to molecular genetic processes. For example, both short (P2O) and long day critical photoperiods are determined by the dynamics of the CONSTANS protein in a range of plants including Arabidopsis [50] and a number of grasses [4], albeit not maize [51]. In rice (*Oryza sativa*), critical short day length has even been successfully predicted from a differential equation model of the diurnal expression patterns of the *CONSTANS* ortholog [52].

Because stability is both important and reasonable to expect given the goals of ecophysiological modeling, it has been argued [5] that finding a putative GSP to be unstable is *prima facie* evidence of a problem. Possible causes of instability include: (1) the model incompletely or incorrectly disentangles G × E; (2) a stable answer exists but the optimizer is insufficiently skilled to find it; (3) undiscovered equifinality is present, and the solutions found depend on low-level algorithmic idiosyncrasies of the optimizer (e.g. method section “equifinality”); and (4) unique best GSP estimates exist that the optimizer can find, but because the model is over-parameterized, the values obtained reflect noise signals that differ between environments.

All sources of instability, whether these or others, are detrimental to the two-step ecophysiological genetic approach to phenotype prediction. Thus, it is critical to know when parameter instability is present, so herein we developed a statistical approach to detect and test for it. The specific question asked was "Do the GSP estimates depend on the particular set of environments used to construct them?" A conceptually simple way to answer this might be to (1) obtain a combination of parameter estimates from one subset of site-years, (2) repeat the estimation with a different subset, and (3) test whether the two sets of parameter estimates differ according to an appropriate statistical test.

A more general and robust approach, however, might be to obtain parameter estimates from many site-year subsets chosen according to a principled method. Preliminary tabulations of the Sobol database revealed that equifinality increased dramatically when fewer than seven site-years were used for estimation (see Results). Therefore, the subset size was set to seven site-years. One method for selection of site-year subsets might be to resample site-years with replacement. However, as shown by analogy in Fig 2b, randomization adds a source of variability to the results that could be of concern given that sampling by replacement would have 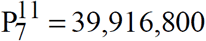 possible site-year subsets. Therefore, analogous to Fig 2b, we used a combinatorics-based sampling pattern leading to more uniformly-distributed site-year subsets by taking all combinations of 11 site-years 7 at a time, of which there are 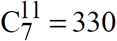 possibilities. To maximize the amount of data available for each line in any subset, we focused on the 539 lines for which observation were available in all 11 site-years.We then conducted 177,870 four-dimensional optimizations to obtain GSP parameter estimates for each of the 539 line × 330 site-year set combinations. These optimizations involved only Sobol database retabulations rather than new model runs, again illustrating the computational efficiency of the database approach. When forced to generate a single result, the database search returned the combination of GSP estimates yielding a minimal RMSE that it happened to encounter first. To focus on the subset that lacked this element of optimizer arbitrariness, we first dropped the 114,314 line × site-year combinations that had ties (i.e. more than one set of GSP estimates yielding the same RMSE). Because our primary interest was in the variability that different site-year combinations might contribute to GSP estimates, we further restricted our attention to the 297 site-year subsets that had at least 100 lines remaining after ties were removed. Each of the 539 lines was present in at least 28 site-year subsets, which was deemed adequate for GSP estimation. These actions left a total of 60,834 estimates for each of the four GSP’s in the study. This became our base group for analysis. We acknowledge that the estimates dropped share a common property (i.e., ties) that might have systematic effects influencing the results. So, in addition to the base group just described above, we also examined the set of (1) all 177,870 GSP sets and (2) the 114,314 results for which ties existed. In both cases we used the optimizer-selected values
We then specified a statistical model to test for stability in parameter estimates across environmental subsets, as follows:

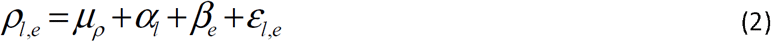
EQN
where *ρ_l,e_* represents an estimate of the GSP *ρ* (i.e. either P1, P2, P2O, or PHINT) for the *l*^th^ line (*l* = 1,2,… 539) obtained from the *e^th^* site-year set (e = 1,2,…297), *μ* is the intercept parameter, acting as an overall mean of GSP *ρ* across all lines and site-year subsets; *α_l_* is the differential random effect of line *l*, assumed to be distributed 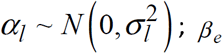 is the differential random effect of the *e^th^* set of site-years, assumed to be distributed 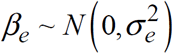 and *ε_l,e_* is the left-over residual unique to the *l*, *e^th^* observed GSP estimate and assumed 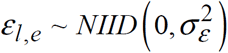. The differential line effects *α_l_* are considered to be random, as is common in field studies of plant population biology. Further, the differential effects of site-year sets, *β_e_*, were treated as random because the corresponding environmental sets are combinations of 7 out of 11 plantings considered to be a representative, if not random, sample of the population of possible site-years to which we are interested in inferring.

If the estimation of any GSP parameter *ρ* were stable across the site-year subsets, one would expect the variance of *β_e_*, namely 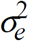, to be zero; alternatively, if estimation is unstable, one would expect 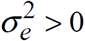. To test this hypothesis set, we fit two competing versions of the statistical model in equation (1), one with and one without the random effect of site-year subsets *β_e_* for each of the GSP’s *ρ* = P1, P2, P2O, and PHINT. For each GSP, we then compared the two competing models using a likelihood ratio test statistic against a central chi-square distribution with half a degree of freedom to account for the fact that the test is being conducted on the boundary of the parameter space. Statistical models were fitted using the liner mixed-effects model package lmer in R [53] with optimization based on the log-likelihood option. The lmer package also calculated the Akaike and Bayesian Information Criteria [AIC [54] and BIC [55], respectively], which allowed for an additional assessment of fit for statistical models that included or excluded the random effects of site-year subsets.

## Results

### Observations vs. Predictions

Fig 3 shows a color-coded scatterplot of observed vs. predicted days to anthesis for 49,491 line × site-year combinations; the cloud of points is concentrated along the identity line, therefore suggesting accurate prediction; the overall estimated RMSE is 2.39 days. Also, there seem to be considerable differences between sites on anthesis days, whereby Florida and Puerto-Rico show very short vegetative durations (ca. 50 d), which are more than doubled in New York (120 d). Empirical correlation coefficients (ȓ) were high across site-years and ranged from 0.86 to 0.95, thus indicating an overall responsiveness across lines to the range of site-year conditions on anthesis dates. The standard deviations of the predicted values and their corresponding observations are 10.336 and 10.639, respectively, which, with the overall empirical correlation coefficient of 0.974, account for a close to 1-to-1 estimated regression slope of observations vs. predictions [i.e. 1.002 = ( 10.639 /10.336) *0.974], as per the established statistical identity between these four sample quantities [56].

**Fig 3.**
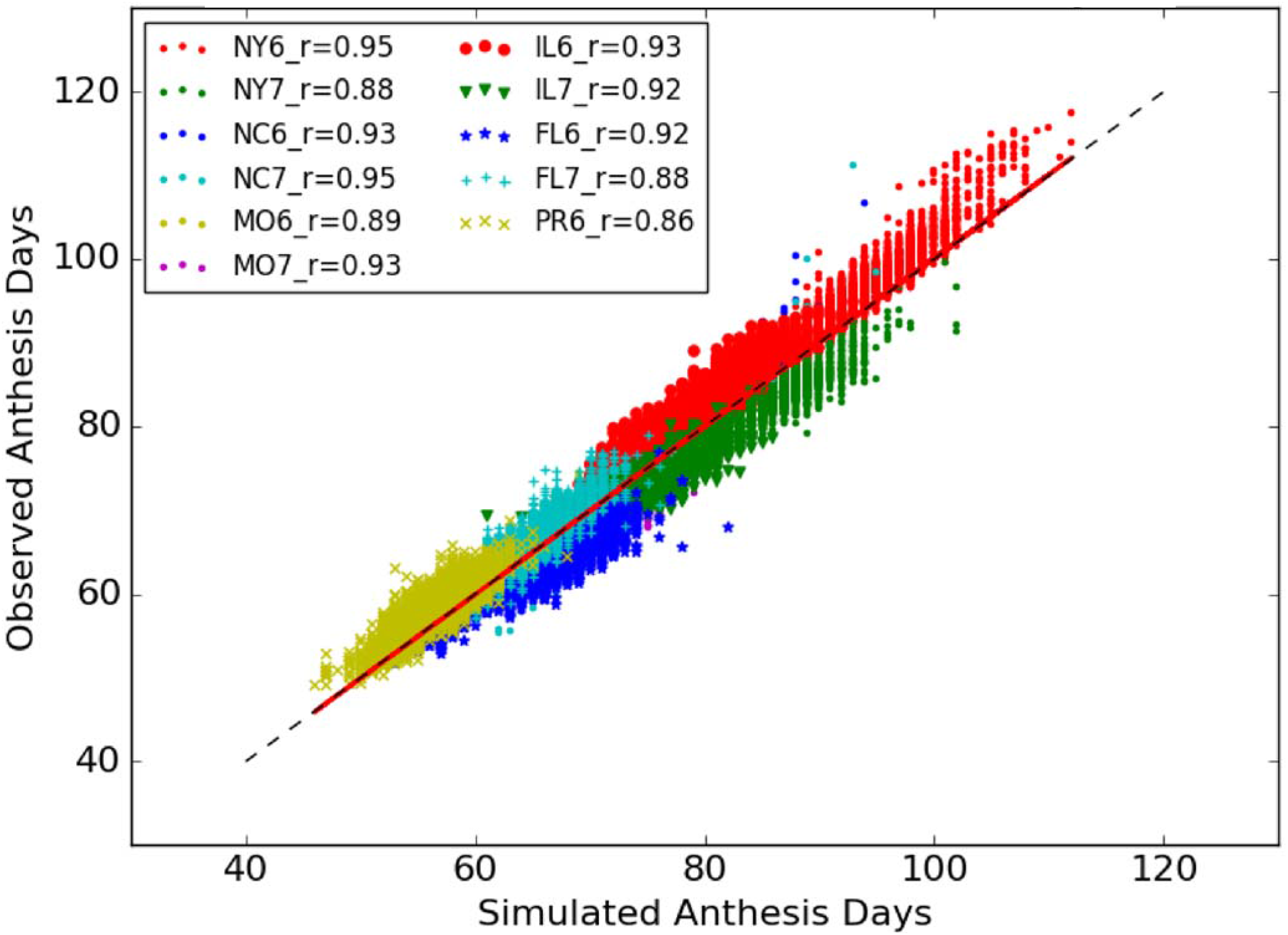
Predicted and Observed anthesis days of all 5,266 lines from 11 site-year combinations. The graph has 49,491 points and an overall RMSE of 2.39 days.

### Equifinality

A more complex picture emerges when the prevalence of equifinality is considered. As noted in 3.4.1, for the 2,254 lines exhibiting equifinality, the number of ties can exceed 1M. The histogram in Fig 4a tabulates the frequency of ties across lines. There are 2,153 lines with fewer than or equal to 40 ties. The line trace along the upper portion of the top and bottom panels shows the average number of site-years in each bin.

**Fig 4.**
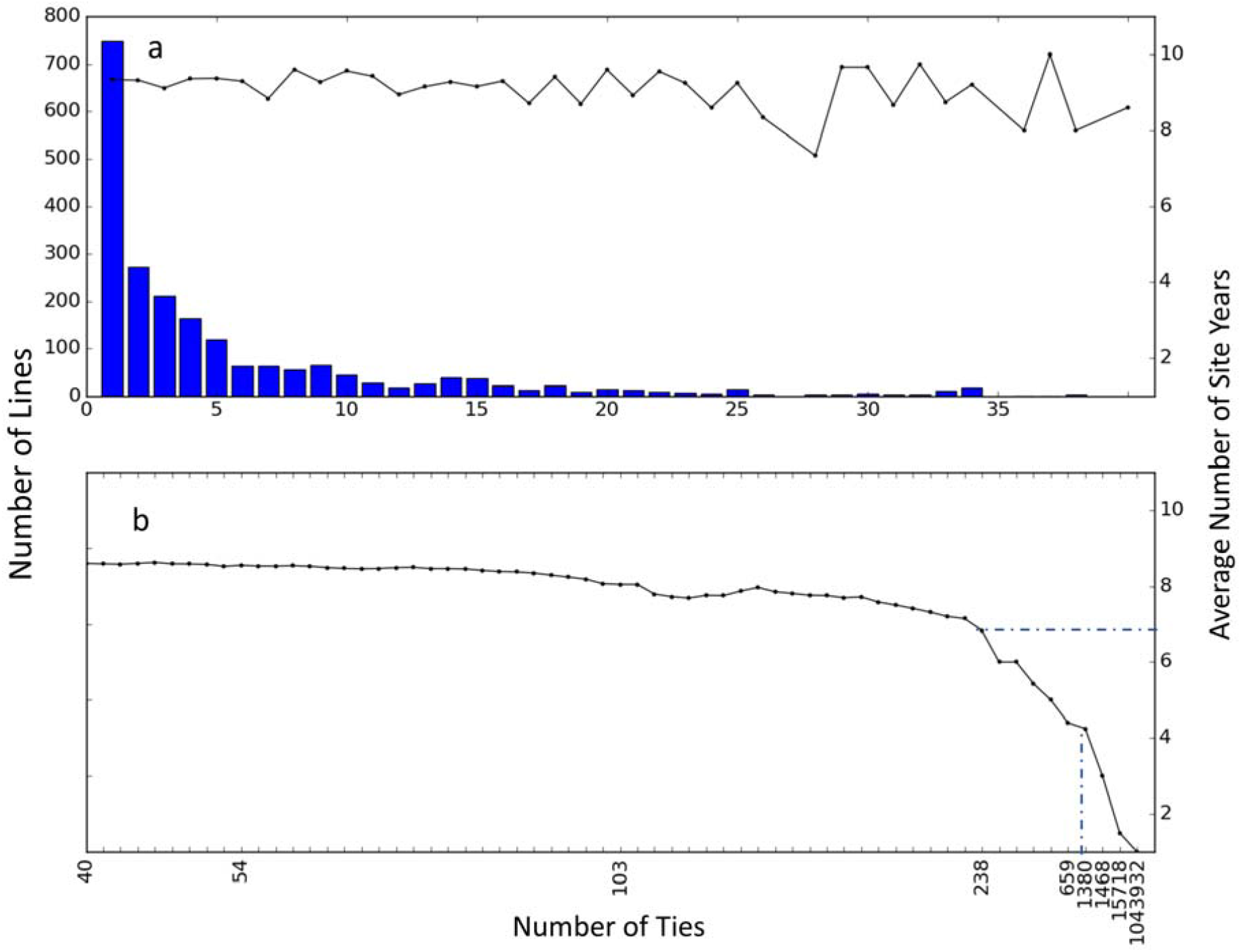
Histogram depicting the frequency distribution of number of ties for 2,254 lines, used here to characterize equifinality. (a) Histogram of number of ties for 2153 lines with fewer than or equal to 40 ties. (b) Continuation of the histogram tail from the upper panel figure representing frequency of ties for the 101 lines with more than 40 ties. The trace at the top of each panel represents the average number of site-year combinations (right axis) used as data for parameter estimation.

In Fig 4a, the empirical distribution of ties was right skewed, thereby indicating that a relatively large number of maize lines had few ties and thus low levels of equifinality. This is particularly true when parameter estimates were computed using data from 7 to 11 site-years (right axis of Fig 4b). Further, the distribution of ties appears to have a very long tail to the right, whereby the number of lines with increasing amounts of equifinality declines very slowly while the number of site-year combinations used for estimation seems to plateau (Fig 4a). This pattern continues into Fig 4b, which shows the 101 lines with more than 40 ties. (No bars are shown in Fig 4b due to scale of the y-axis, as each bin generally contains one to three lines.) Interestingly, the number of ties, and thus equifinality, seems to increase precipitously for the 56 out of 5,266 lines that have fewer than seven site-years of data (Fig 4b).

As the number of ties increases, one can expect that the range of indistinguishable estimates for any GSP will widen. To illustrate this phenomenon, a set of GSP estimates were obtained using just two illustrative site-years (NY6 and NY7) so as to artificially inflate equifinality. Fig 5 shows scatterplots of coordinate pairs of either predicted (a) or observed (b) values for anthesis days from NY6 (horizontal axes) and NY7 (vertical axes). Points in each scatterplot are color-coded to represent the number (on a log_10_ scale) of tied GSP combinations. Each tied GSP combination, when simulated using the weather data for NY6 and NY7, predicts the same anthesis dates that form the point’s coordinates. Dark red indicates 235,976 ties and blue indicates 1 tie. It is reasonable to expect that as the number of ties increases, the range (max minus min) of the equifinal estimates will increase. The size of each circle indicates the range of tied P1 estimates expressed as a percentage of the mean. These percentages extend from 0.36% to 65.68%. The association of redder colors with larger circles indicates that estimate ranges do, indeed, increase with the level of equifinality.

**Fig 5.**
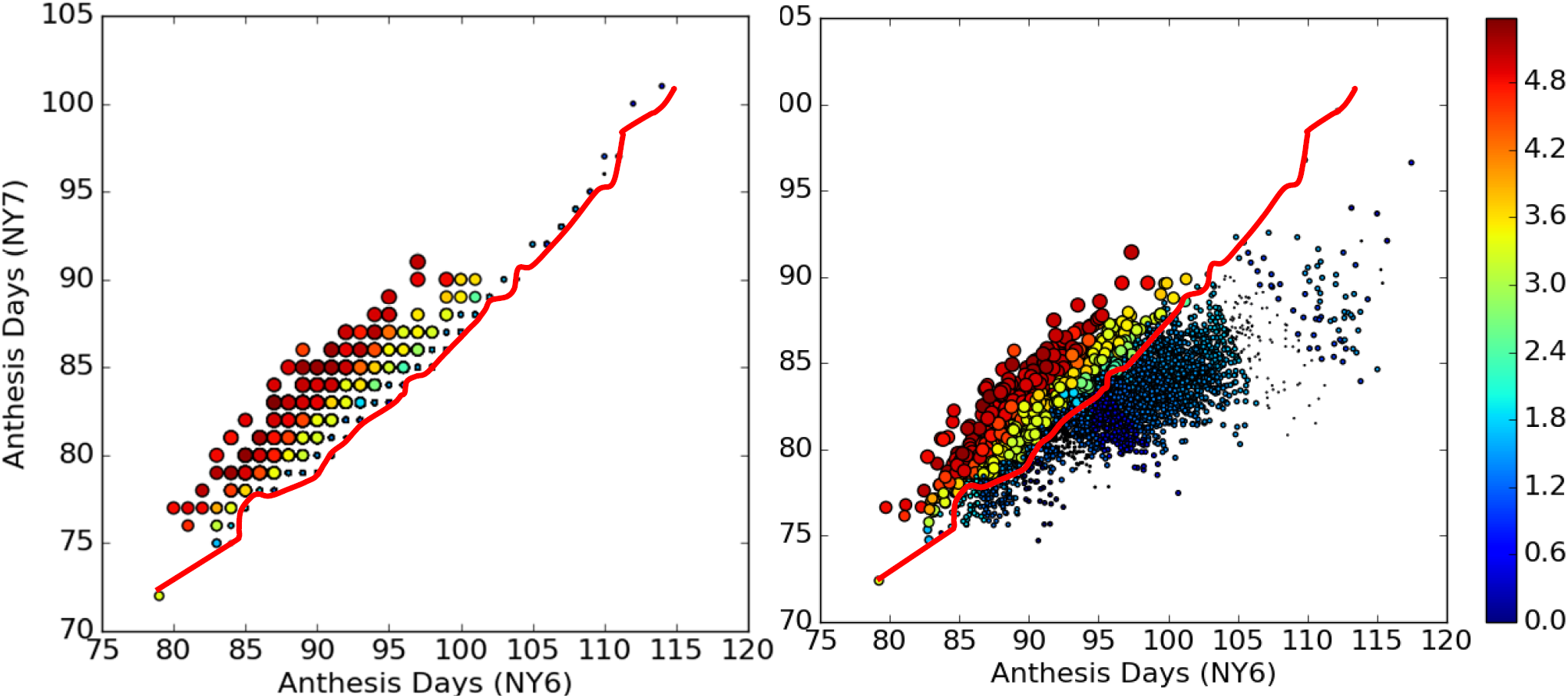
Phenotype space plots of predicted (a) and observed (b) values of anthesis dates for site-years NY6 and NY7. The marker sizes and colors respectively express the levels of equifinality based on number of ties for P1 (log_10_ scale) and the relative ranges of its tied values. The red line is explained in the text.

This is an example of a phenotype space plot that can be used to show how properties of interest (e.g. number of ties and estimate ranges in this case) are distributed across the range of predictions made by the model given the weather in a pair of site-years. Notice that (1) the cloud of observed points (Fig 5b) is more dispersed than that of the predicted points (Fig 5a), suggesting that model responses to the environment were less plastic than those of real plants and (2), as indicated by the red lines, the lowest numbers of ties in Fig 5b (blue points) appear to fall in empty regions of Fig 5a where predictions are lacking. This pattern has important consequences to be explained later in section “model expressivity”.

### Interrelationships between parameter estimates

Fig 6 presents a combined plot depicting histograms of GSP parameter estimates based on all 5,266 lines along the main diagonal and corresponding pairwise GSP scatterplots in the upper right panels. The GSP estimates were obtained using all site-years. The lower left panels in Fig 6 show the estimated Pearson correlation coefficients 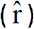, estimated regression slopes 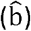, and corresponding *p*-values for each mirrored scatterplot. Two immediately apparent features on the scatterplots are to be noted, which might readily escape notice in data sets with fewer lines. The first is the pronounced banding pattern appearing in all plots except, perhaps, P2O vs. PHINT. Most bands seem to be linear except for those on the scatterplot of P2O and P2 plot, which exhibits curvilinearity. The second is the pronounced vertical gap in all P2O scatterplots. In an attempt to understand the reasons for such patterns, the authors explored multiple seemingly plausible hypotheses, ranging from genetics to input file coding quirks (e.g., unintended rounding of parameter values) and many more, all of which were tested and discarded. Ultimately, the results presented in the following sections provided the explanations.

**Fig 6.**
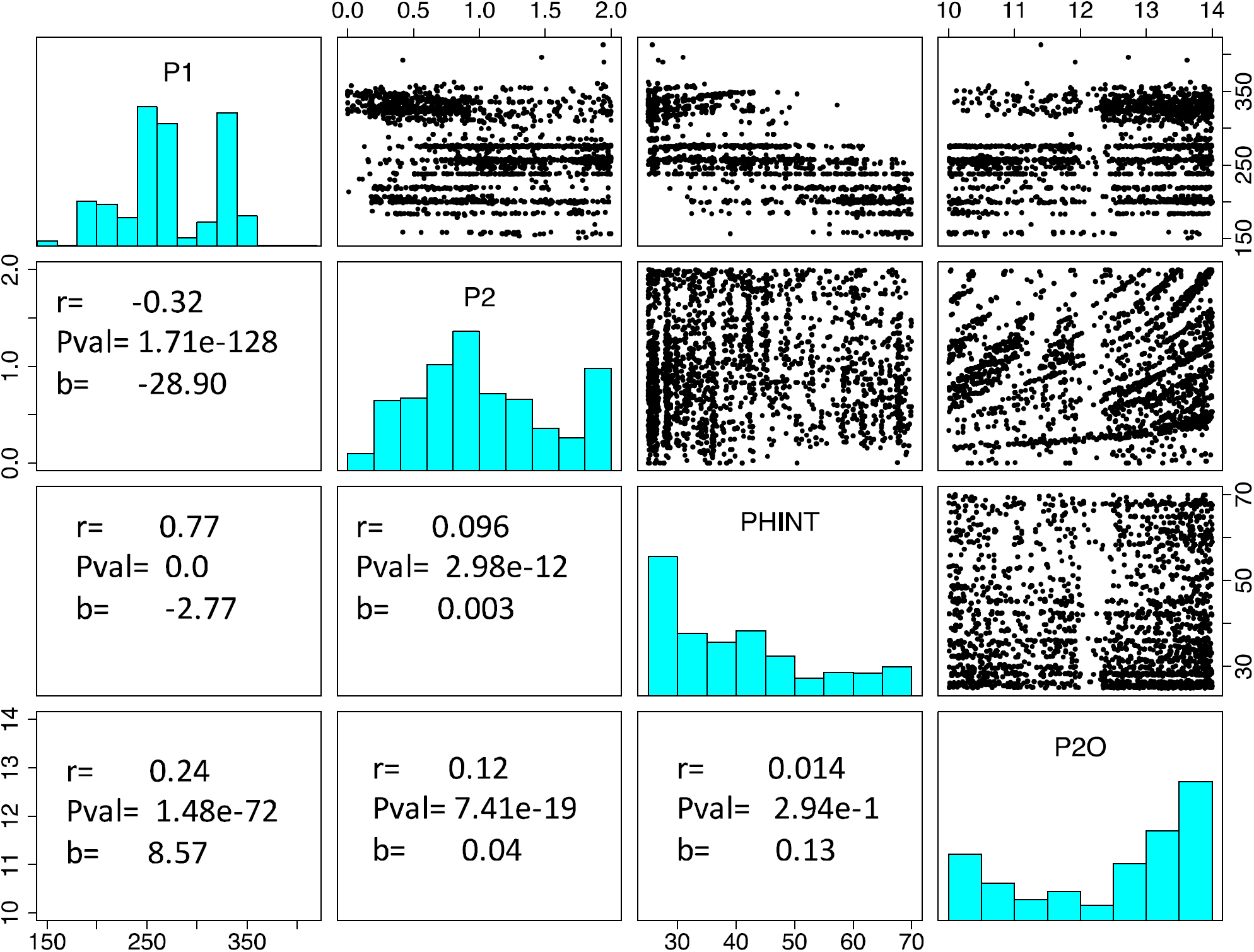
Empirical distribution of selected GSP parameter estimates (main diagonal), pairwise scatterplots (upper right triangle) and empirical estimates of Pearson correlation coefficients, regression coefficients and p-values (Lower left triangle). Each dot in the scatter plots represents a pair of GSP estimates from a single line.

### Model expressivity

The first clue to the cause of the banding pattern emerges from the phenotype space plots in Fig 7. Each plot corresponds to an independent fit to just one particular pair of site years. The blue regions in each panel of Fig 7 outline predicted anthesis date pairs for two consecutive years in a given site, where model prediction are constrained by the bounds imposed on the range of values allowed for each of the four GSP’s (Table 2). Also, for each panel in Fig 7, a dot depicts an observed anthesis date pair for a line present in a given site in both 2006 and 2007. Yellow (red) dots represent observed anthesis date pairs that the model was able (unable) to reproduce. We characterize each observation corresponding to a yellow (red) dot as “expressible” (“inexpressible”). Except for the two North Carolina site-years, there were many lines (Table 3) for which observations on anthesis date could not be predicted despite: (1) the seeming breadth of GSP values allowed by Table 2; and (2) the fact that the model was only being asked to match two data points, which would seem to greatly relax the constraints on GSP estimates.

**Fig 7.**
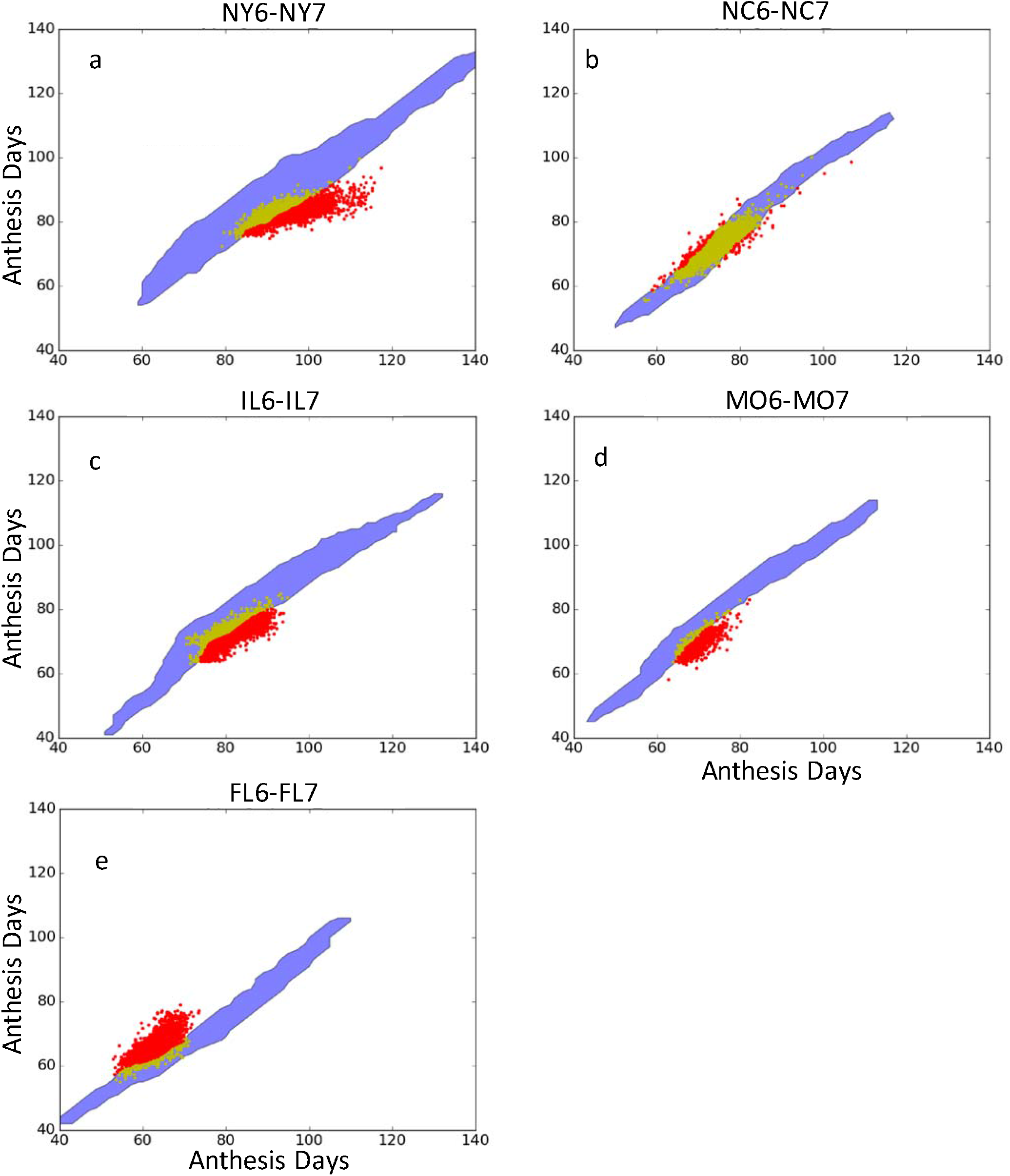
Phenotype space plots for predicted and observed anthesis dates. Each panel corresponds to a pair of site-years for which fits were done. Regional color codes are described in the text.

**Table 3.** Numbers of model expressible and inexpressible observations for selected site-year pairs.

| Lines that are<sup>a</sup>: | NY6/NY7 | NC6/NC7 | IL6/IL7 | MO6/MO7 | FL6/FL7 |
| --- | --- | --- | --- | --- | --- |
| Expressible | 2189 | 4964 | 2024 | 146 | 193 |
| Inexpressible | 2542 | 168 | 2946 | 637 | 3339 |

<sup>a</sup> These numbers refer to lines with data in both years of each pair and therefore do not precisely align with Table 1.

This begs the question as to what would happen to model expressivity if an even broader range of GSP values were allowed. In an attempt to investigate in a computationally efficient way how the outputs of a more conventional optimizer might appear when viewed in phenotype space, the CERES-Maize anthesis date routine was ported to Python and fit to NY6/NY7 via Differential Evolution (DE) [57]. DE is a well-established (63K Google Scholar hits on “Differential Evolution” as of October 21, 2016) and highly effective evolutionary algorithm that embodies mechanisms reminiscent of techniques ranging from the Nelder-Mead Simplex [58] method to Particle Swarm Optimization [59]. Among the algorithm’s initiating inputs is the range of parameter values within which to search, which were set as shown in Table 4. These ranges are greatly broadened from that used in the database search (Table 2); in fact, the values in Table 4 are intentionally broader than biological experience would suggest as reasonable.

**Table 4.** Extended range of parameter values used for DE search.

| Parameter | Definition | Unit | Min | Max | Percent of<br>Sobol Range |
| --- | --- | --- | --- | --- | --- |
| P1 | Thermal time from seedling emergence to<br>end of juvenile phase | GDD (°C) | 75 | 600 | 175% |
| P2O | Critical photoperiod hour | hrs. | 6 | 21 | 300% |
| P2 | Days of anthesis date delay for each hour by<br>which the day length exceeds P2O | rate | 0 | 6 | 375% |
| PHINT | Phylochron interval (Interval between<br>successive leaf tip appearances) | GDD (°C) | 20 | 110 | 200% |

Fig 8 shows overlapping predictions based on the database search under the range of parameters in Table 2 and on the DE search under the extended range of parameter values (Table 4). Specifically, the light blue area represents the anthesis date region that was reachable through predictions based on the database search. In contrast, the dark blue area is the predicted anthesis date region within which the DE algorithm converged. Note the almost perfect overlap of the lower edges of the light blue (i.e. database search) and dark blue (i.e. DE search) areas, indicating that, despite its much larger starting parameter search space, DE did not extend model predictions. This suggests limitations in model expressivity that go beyond the method of parameter estimation or the initial parameter space used for the search.

**Fig 8.**
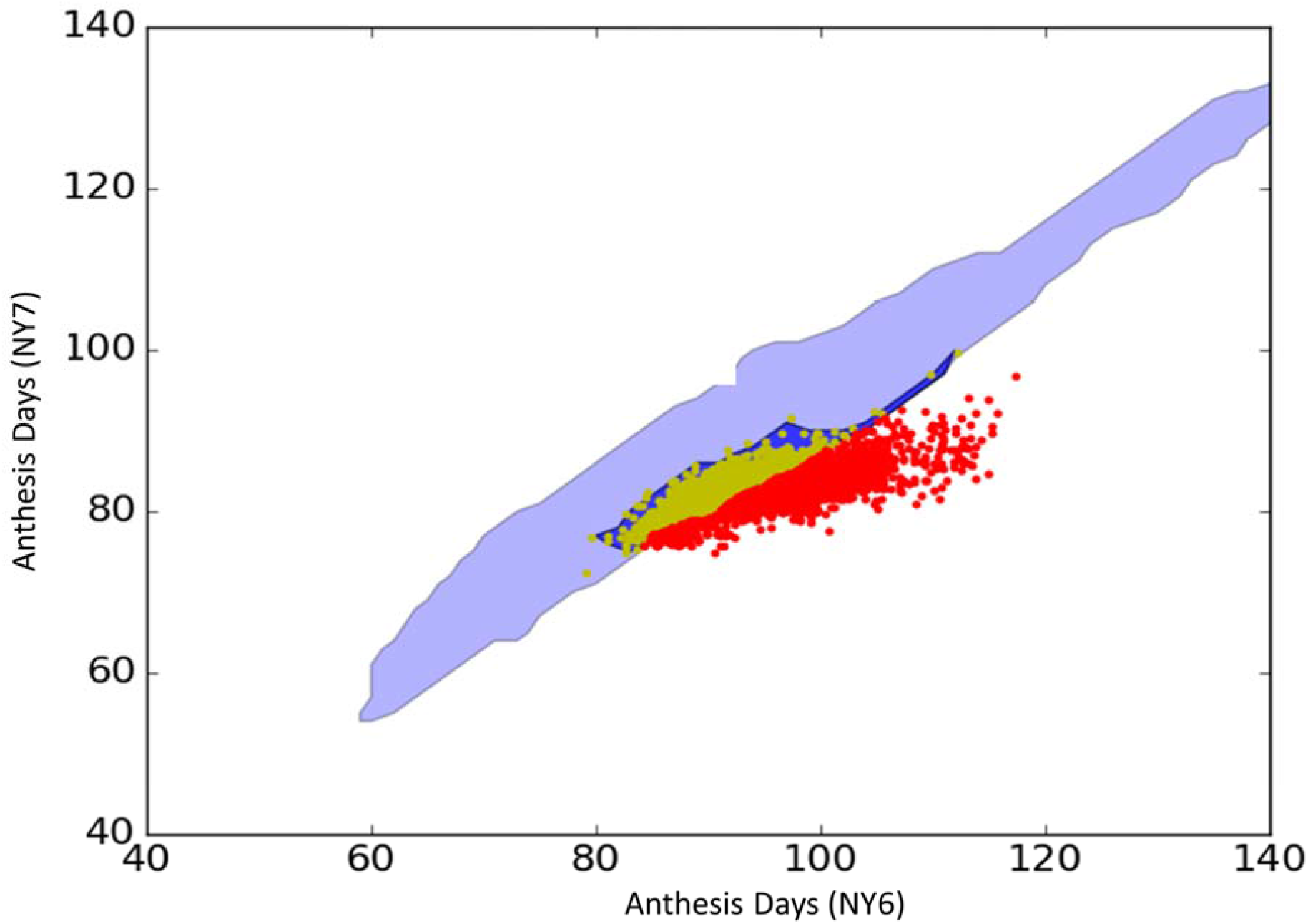
Superimposed anthesis date results using NY6 and NY7. Data illustrating that searches via database and DE optimization over a much larger parameter space are equally unable to reproduce the observations for lines shown as red dots.

As a corollary, it is worth noting that more site-years of data of similar quality are unlikely to improve model expressivity, as illustrated by the following thought experiment. Suppose a community has developed the univariate deterministic model *y* = arctan(*θ*), where *θ* is a parameter, with 0 ≤ *θ* ≤ 10 by solid prior knowledge and y is some dependent variable of interest. Assume that this is viewed as a very complex model requiring simulation to solve. The community understands that no model is perfect, but no specific flaws of this one are known. Extant data for y ranges from 1.31 to 1.61 and yields the point estimate 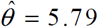 (RMSE = 0.12). Due to its complexity, no one has noticed that the model cannot reproduce any ***y*** > arctan(10) =1.47 or, for any *θ*, a *y* > *π*/*2* ≈ 1.57. Now suppose that: a very large set of new *y* data is collected. Depending on the distribution of the new data either: (1) a new 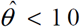 will be found or (2) 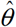 will rise significantly above 10, leading to a rejection of the model. However, what will not happen is that the increase in data will enable observations >1.57 to be reproduced. The model simply lacks the expressivity to do so. Analogously, increasing the amount of anthesis date data may narrow GSP estimate confidence limits, but the reachable region of predicted phenotype space is unlikely to extend beyond the edges of the light blue regions. Therefore, any improvement in the ability to predict the large numbers of red points in Figs 7 and 8 is unlikely.

Given these issues, a sensible follow-up question might be about what specific GSP estimates were reported for the red points? Here we report answers only for P1.

Fig 9 shows scatterplots of P1 and P2O estimates generated using data from NY6 and NY7 via the database search and DE. The color coding is consistent with that in Fig 7a. The pronounced bands at ca. P1=250 in both panels are immediately striking - although the scale is small, a corresponding band is quite evident at the same position in Fig 6. A tabulation reveals that, of all 4,731 lines represented in the Fig 9a, 3,227 (68.2%) have estimates of P1 ranging from 245 to 260. Of these, 1,493 are expressible (yellow) and 1,734 (red) are not expressible. Out of the total 4,731 points in the graph 2,189 (46.2%) are expressible and 2542 (53.8%) not. The Fig 9b has similar proportions of expressible and inexpressible points (2327, 49.1%; and 2404, 50.9%; respectively), reinforcing the similarity of results for parameter estimates from DE and database searches. The differences are likely due to the ability of DE to explore the parameter space continuously whereas the database search is restricted to the predefined discrete Sobol points. Still, one may wonder why so many P1 estimates are near the 250 degree-days? Fig 10 reveals the answer.

**Fig 9.**
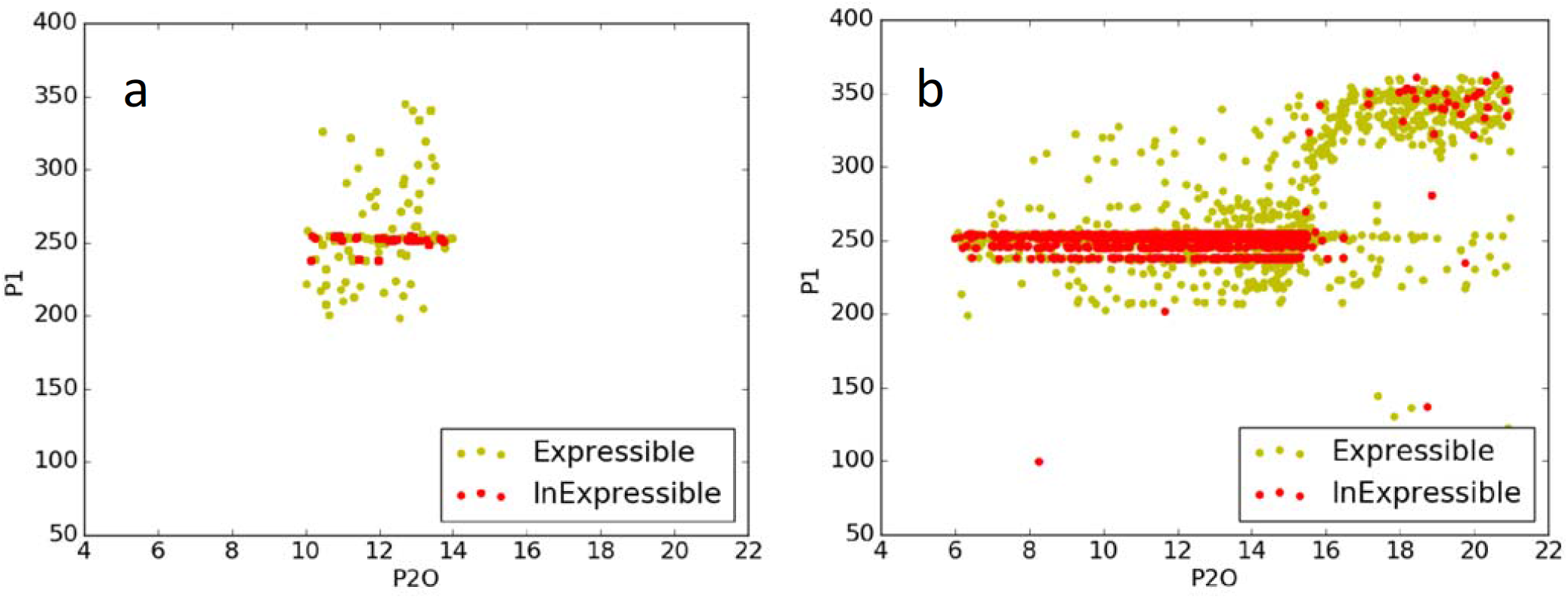
Scatterplot of P1 vs. P2O estimates using data from NY6 and NY7 based on the database search (a) and Differential Evolution (b). Yellow and red dots are, respectively, observations characterized as expressible and inexpressible by model predictions.

**Fig 10.**
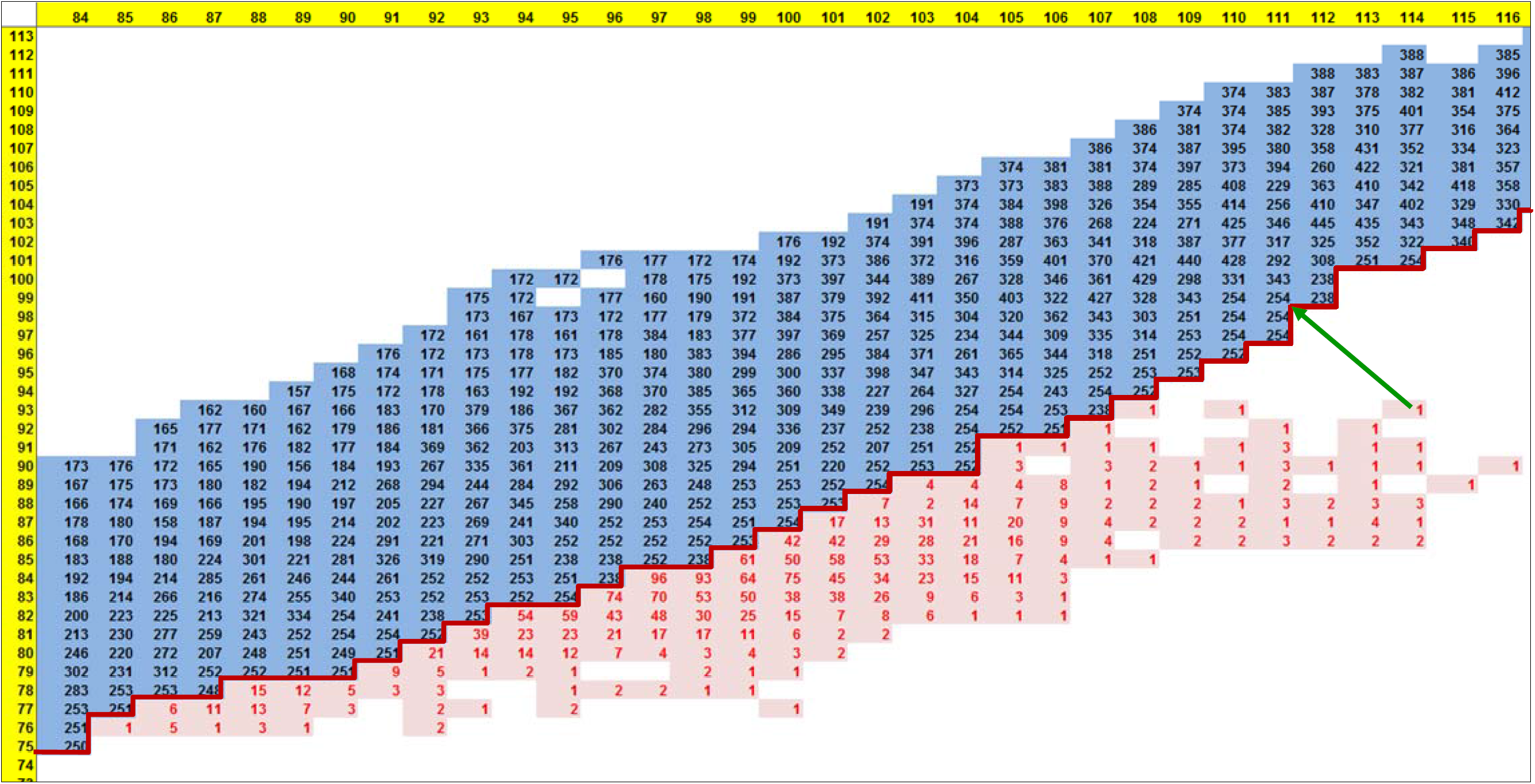
P1 estimates from the database search (black) and the numbers of lines with inexpressible observations (red). Figure arranged in a tableau organized as a phenotype space plot corresponding to the center portion of Fig 8. The dark red line is the expressibility frontier and the green arrow shows the P1 value (254) from the GSP combination that minimizes the RMSE for one illustrative line. Horizontal and vertical yellow strips are the anthesis dates for NY6 and NY7

The numbers in black are the “first-best-found” P1 estimated values that generate the corresponding row × column anthesis date combinations. A comparison with the corresponding dot colors and sizes in Fig 5b indicates that, on the frontier (red borders Figs 5a,b and 10) between expressible and inexpressible observations, there was essentially no equifinality and, concomitantly, narrow ranges of P1 values. Fig 10 shows that the P1 values along the frontier were all quite close to 250. For lines with observations falling outside the frontier, the RMSE was minimized by assigning GSP values associated with the closest achievable dates, i.e. those directly on the frontier. Therefore, all the lines counted by the red numbers were assigned P1 values that are very close to 250 and have essentially no equifinality. The green arrow in Fig 10 illustrates this phenomenon for one line. The nearest P1 estimate is 254 and the length of the arrow (ca. 5.8 days) is proportional to that line’s RMSE. Specifically, in this case the length is 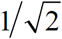 times the RMSE because there are *n* = 2 site-years.

Recall that the upper limit placed on P1 was 450 (and 600 in the DE search), therefore this outcome is likely not an artifact of constraints in the GSP search space but, rather, a result of poor model expressivity, that is the model inability to predict anthesis date pairs beyond those on the frontier. This mechanism accounts for the P1 band at 250 in Fig 9a. Furthermore, as previously presented, more data cannot improve the prediction of inexpressible lines, the banding in Fig 6 is not surprising.

### P2O gap

We now investigate the vertical gap in scatterplots involving P2O estimates (Fig 6), which documents the intricacy of the interactions that can occur between model mechanisms, parameter ranges searched, optimization algorithms used, and environments included. Exploratory re-tabulations of the Sobol-based parameter database revealed that the P2O gap was clearly present in the three site-years having shorter day lengths (FL6, FL7, and PR6 but absent in fits obtained by only including the remaining eight site-years with longer days (Fig 11). Fig 12 shows that a substantial number of observations for short-day site-years are outside the predicted phenotype ranges expressible by the model under either database or DE optimization. As described in section “CERES-Maize model”, the model operated by calculating the number of leaves initiated by the end of Stage 2 and predicts anthesis only after leaves are fully emerged. For any line, leaf number was a constant across all site-years, namely P1/(2×PHINT)+5. The variation of anthesis dates across plantings was such that there were few, if any, combinations of P1 and PHINT that were compatible with the data from all site-years. Therefore, the optimizer relied more heavily on the P2 and P2O parameters.

**Fig 11.**
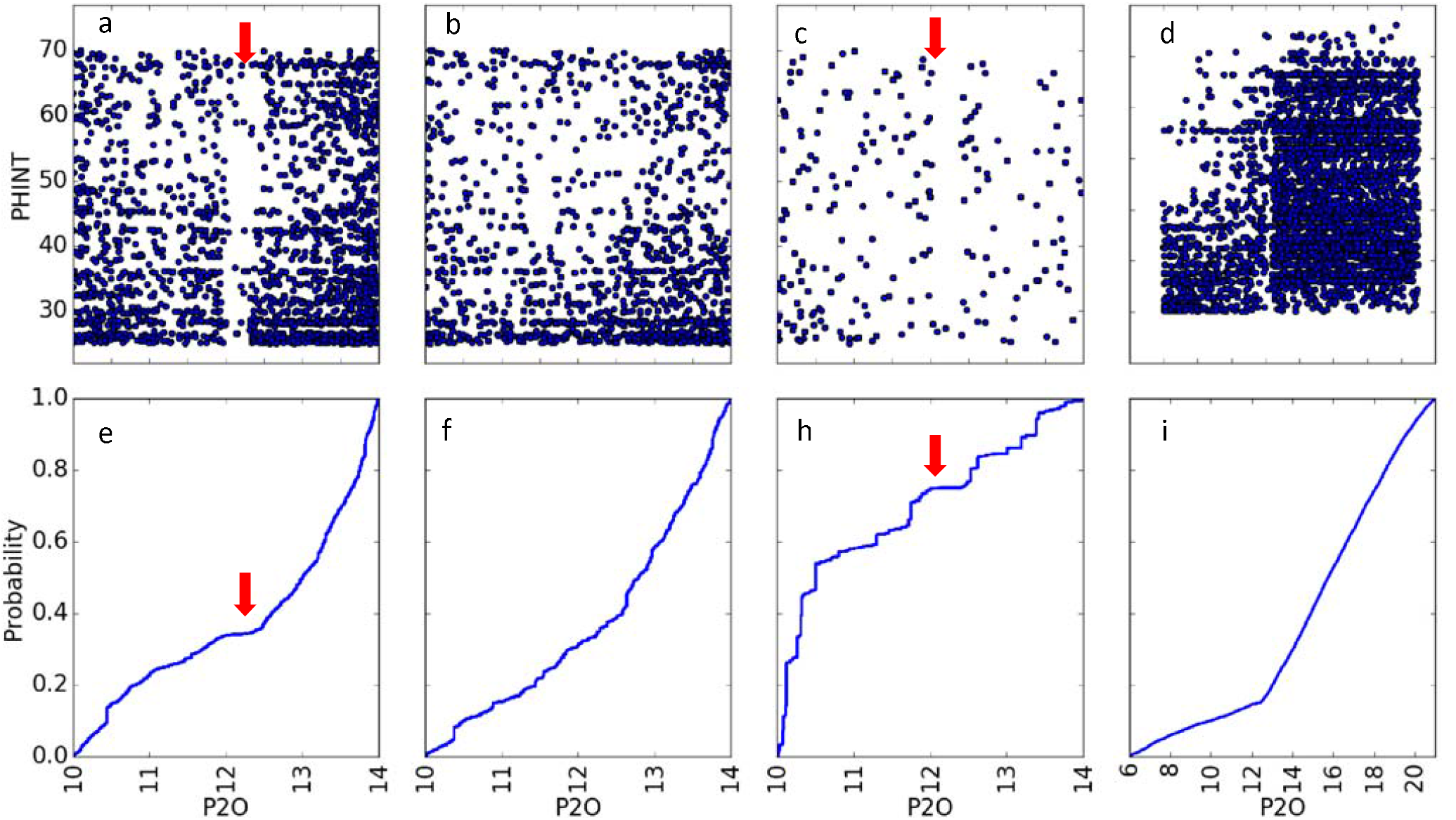
P2O and PHINT scatter plots (top row) and P2O cumulative density functions (bottom row). (a & e) all 11 site-years, ( b & f) longer day site-years, (c & g) shorter day site-years based on the database approach, and (d & i) shorter day site-years using the DE approach. All horizontal axes in both rows have the same scale.

**Fig 12.**
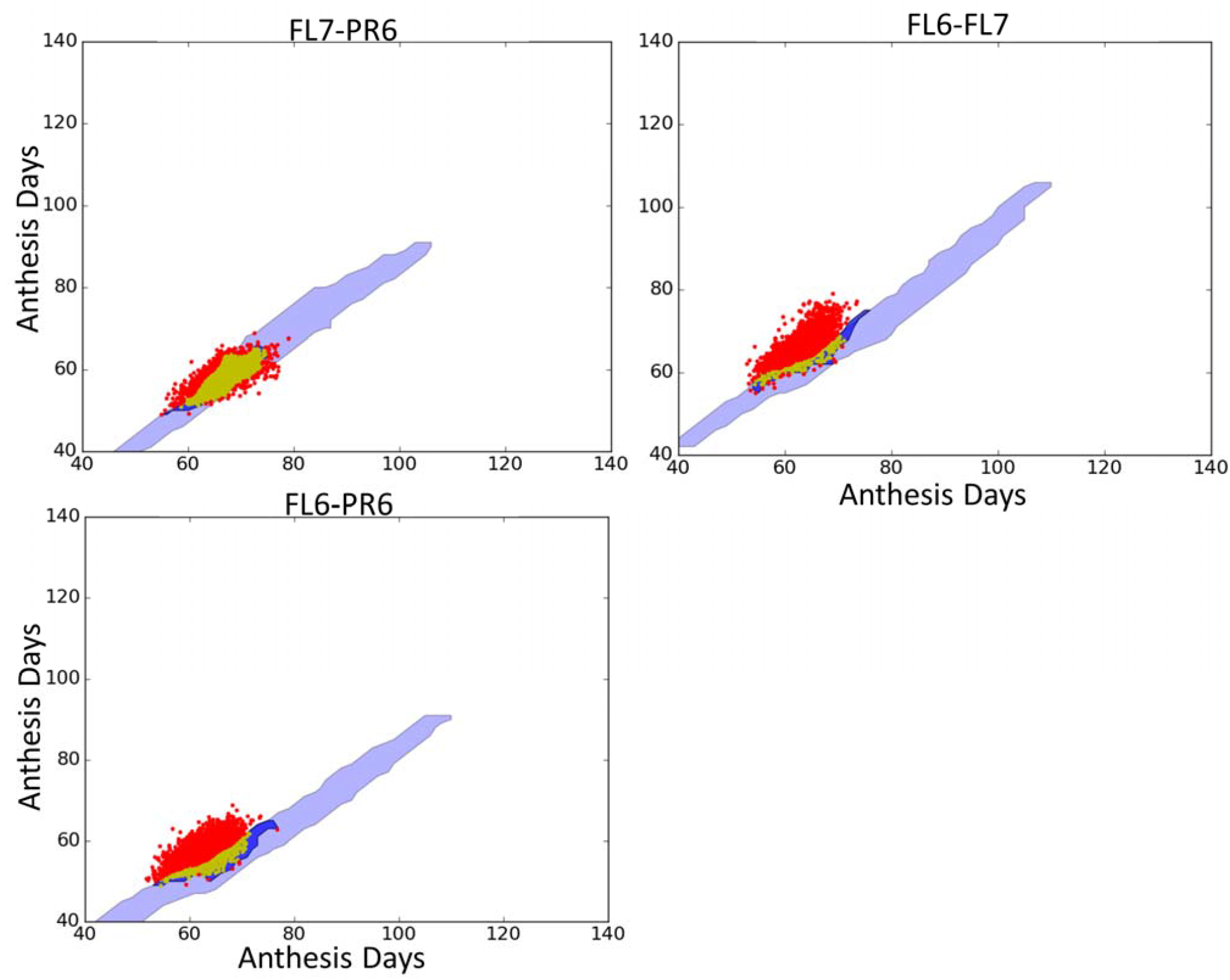
Phenotype space plots of observed and predicted values based on the three site-years with shorter days. Note the large number of points in the FL6-PR6 and FL6-FL7 plots that lie above the dark blue prediction region based on DE.

Specifically, the optimizer settled on very small P2O estimates, much smaller than the short southern photoperiods. Instead, the optimizer relied on P2 estimates to generate anthesis date predictions that were delayed to the greatest extent possible by lengthening Stage 2. Recall that P2O values above the day length make Stage 2 only four days long, which is not enough time for temperature differences to accumulate the needed variation. The abundance of low P2O estimates thus created the gap observed in scatterplots of P2O with other GSPs (Fig 11a). In contrast, the photoperiods in the remaining longer-day site-years exceeded the maximum allowed P2O values in the P2O database search during (and long after) the juvenile period. Therefore, there was no empty band in the scatter plot (Fig 11b) because the optimizer was able to exploit delays for any value of P20.

With the broader range of parameter values available to the DE runs and the increased flexibility available between P1 and PHINT, other options became available. In particular, in many cases DE found GSP combinations wherein P2O exceeded the southern day lengths so photoperiod had no influence on anthesis date and no gap artifact was generated (Figs 11d,i). P1 and PHINT thus became the major explanatory parameters. This is shown in Fig 13, whereby for each line, the parameter differences are plotted against the RMSE differences that result from changing the estimation methods from database to DE optimization. The DE estimate of P2O were larger in 4,507 out of 5,240 lines (87%; Fig 13d), almost always by enough to put it above the local day lengths. In tandem, P1 values fell in 3,559 lines (Fig 13a), whereas PHINT rose in 4,102 lines (Fig 13c).

**Fig 13.**
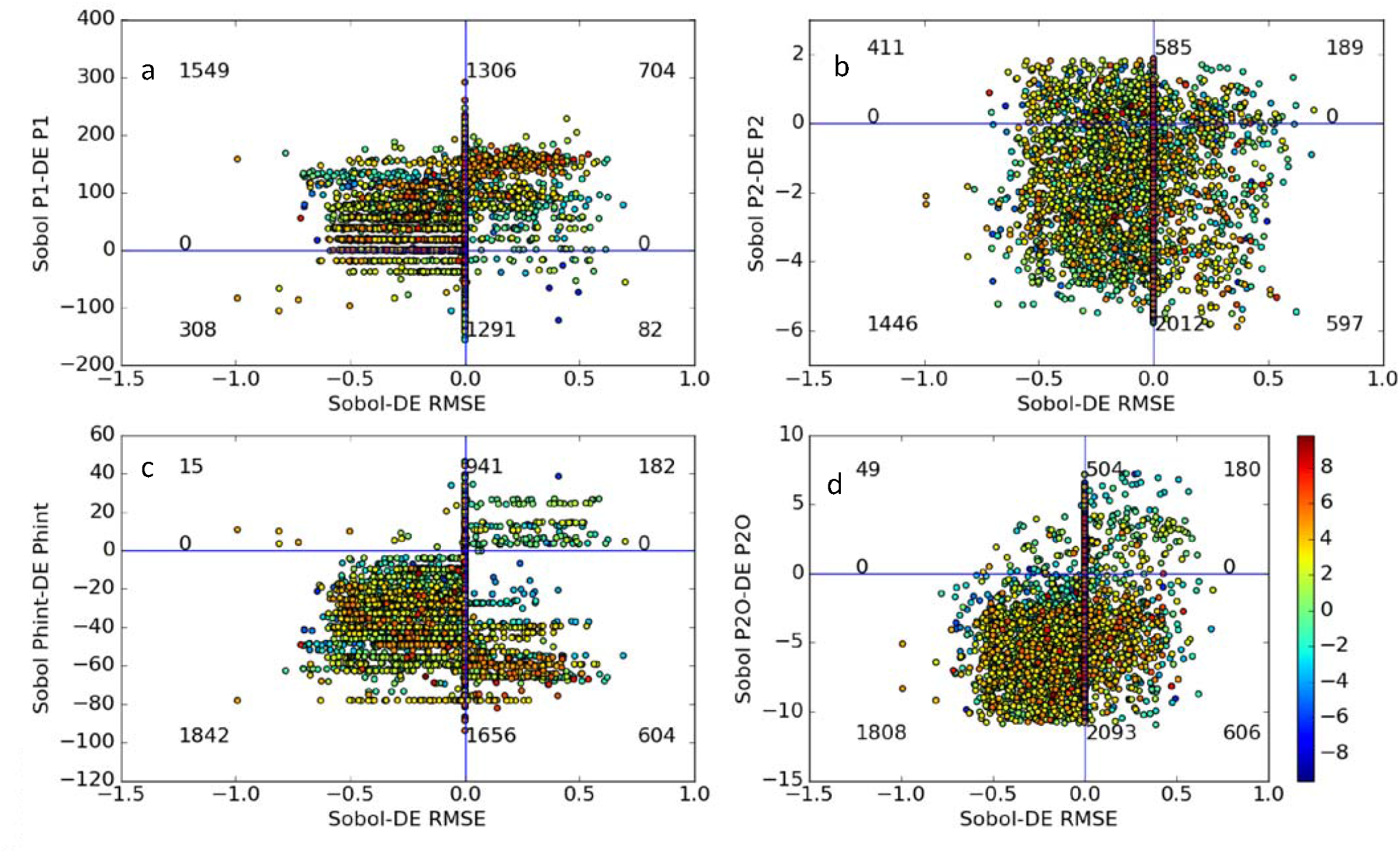
The differences in parameter estimates from database search vs. DE (vertical axes). Figure plotted against the corresponding difference in RMSE for 5240 lines in FL6, FL7, and/or PR6. The color encodes the sum of residual (observed minus mean) across site-years for each line.

Note, however, that for *any* (P1, PHINT) combination, *any* P2O that exceeds the local day length will give the same RMSE - a clear source of equifinality. Thus, the changes in P2O will not, in all likelihood, lead to values that can be more closely related to genetics. Moreover, because of the limits on model expressivity, none of the DE solutions gave significantly better fits than the database estimates. This is why virtually all points in Fig 13 had DE RMSE’s within 0.5 days (horizontal axes) of the database-based parameter estimates. This, too, is an illustration of equifinality because the two optimizers were finding different GSP estimates although the RMSE were of similar magnitude.

### Tests for stability of GSP estimates

Table 5a shows the effect of including or excluding the effect of different subsets of site-years on the modeling of estimates (Equation 1) for each GSP for the base set. For all GSP parameters, AIC and BIC values were considerably smaller for models that included the random effect of site-year subsets, *β_e_*, therefore suggesting non-negligible variability across site-year subsets on the GSP estimates. The table illustrates the size of the site-year set effects as follows. For scaling purposes, we provide the estimated intercept, 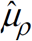, which also serves also as an estimated GSP grand mean across all lines and site-year subsets. The Index of Variability (expressed as a percent) is the standard deviation of the *β_e_* effect normalized by the grand mean. The percentage of the total GSP variance 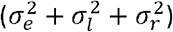 attributable to site-year subsets is also shown. Both of these descriptors indicate substantial variability between site-year sets, with indexes of variability ranging from 5.9% for P2O to 33.6% for P2 and over 20% of the total variance related to site-year sets for all GSP’s.

**Table 5a.** Estimated log likelihood, fit statistics, selected summary measures, and a likelihood ratio test for competing statistical models fitted on GSP estimates with and without the random effect of site-year subset, based on GSP estimates for the base group (N=60,834).

| GSP | Log<br>likelihood<br>w/o (top)<br>and w/ (bot)<br>a site-year<br>set effect<sup>a</sup> | AIC w/o (top)<br>and w/ (bot) a<br>site-year set<br>effect<sup>b</sup> | BIC w/o (top)<br>and w/ (bot)<br>a site-year<br>set effect<sup>b</sup> | GSP Grand<br>Mean<br>$\hat{\mu}_p$ | Index of<br>Variability<sup>c</sup><br>$\sigma_e / \hat{\mu}_p$ | Variance<br>pcts. for<br>site-year<br>sets<sup>c</sup><br>$\sigma_e^2 / \sigma_{tot}^2$ | Chi-square<br>test<br>statistic | Chi-square<br>*p*-value<sup>d</sup><br>(df =0.5) |
| --- | --- | --- | --- | --- | --- | --- | --- | --- |
| P1 | -338046<br>-322689 | 676098<br>645386 | 676125<br>645422 | 264.625 | 12.30 | 34.38 | 30714 | 10<sup>-13334</sup> |
| P2 | -46154<br>-25237 | 92313<br>50482 | 92340<br>50518 | 1.037 | 33.55 | 33.92 | 41833 | 10<sup>-18163</sup> |
| P2O | -105304<br>-95357 | 210614<br>190723 | 210642<br>190759 | 12.2440 | 5.88 | 27.83 | 19894 | 10<sup>-8635</sup> |
| PHINT | -254875<br>-246903 | 509756<br>493815 | 509783<br>493851 | 44.167 | 15.44 | 22.62 | 15943 | 10<sup>-6919</sup> |

<sup>a</sup> Larger is better <sup>b</sup> Smaller is better <sup>c</sup> Chernoff upper bound on Chi-squared cum. dist. function.

The Chi square values from the likelihood ratio test and the associated p-values are presented in the last two columns of Table 5a. The extreme p-values demonstrate that the GSP values depend on the set of site-years used to estimate them. Therefore, the GSP’s are not, in fact, genotype specific despite the goodness-of-fit displayed in Fig 3. This result is completely understandable given the range of artifacts due to equifinality and model expressivity issues identified above.

Table 5b shows the results when only estimates having ties are tested (left) vs. an analysis that includes all estimates (right). The former corresponds to estimates for lines whose observations fall inside the expressivity frontier and the latter includes the estimates for all lines. It is clear that the grand means, index of variability, and percentages of GSP variance are highly similar between all three groupings in Table 5. Also, all p-values are extremely significant and increase with the amount of data used.

**Table 5b.** Summary measures and likelihood ratio p-values for competing statistical models fitted on GSP estimates with and without the random effect of site-year subset from data only having ties (left) and all data (right). Larger is better Smaller is better Chernoff upper bound on Chi-squared cum. dist. function.

| GSP | GSP<br>Grand<br>Mean<br>$\hat{\mu}_p$ | Index of<br>Variability<sup>c</sup><br>$\sigma_e / \hat{\mu}_p$ | Variance<br>pcts. for<br>site-year<br>sets<sup>c</sup><br>$\sigma_e^2 / \sigma_{tot}^2$ | Chi-square<br>$p$-value<sup>d</sup><br>(df =0.5) | GSP<br>Grand<br>Mean<br>$\hat{\mu}_p$ | Index of<br>Variability<sup>c</sup><br>$\sigma_e / \hat{\mu}_p$ | Variance<br>pcts. for<br>site-year<br>sets<sup>c</sup><br>$\sigma_e^2 / \sigma_{tot}^2$ | Chi-square<br>$p$-value<sup>d</sup><br>(df =0.5) |
| --- | --- | --- | --- | --- | --- | --- | --- | --- |
| With Ties (N=114,314) |  |  |  |  | With all Data (N=177,870) |  |  |  |
| P1 | 273.5 | 11.37 | 29.77 | $10^{-23283}$ | 270 | 11.48 | 29.94 | $10^{-34955}$ |
| P2 | 0.9137 | 36.33 | 35.23 | $10^{-34723}$ | 0.9593 | 35.5 | 33.8 | $10^{-52518}$ |
| P2O | 12.49 | 4.43 | 19.70 | $10^{-11883}$ | 12.42 | 4.88 | 21.27 | $10^{-19806}$ |
| PHINT | 43.57 | 18.65 | 26.31 | $10^{-17348}$ | 43.94 | 17.3 | 24.35 | $10^{-23740}$ |

<sup>a</sup> Larger is better <sup>b</sup> Smaller is better <sup>c</sup> Chernoff upper bound on Chi-squared cum. dist. function.

## Discussion

Since their inception, ecophysiological models have been evaluated in terms of predictive ability, which are superb in many circumstances [60]. The model parameters were considered to be *inputs* whose genesis was secondary as long as the model outputs proved useful. However, as often happens in science, perceived needs, desiderata, and requirements escalate as technologies evolves. In particular, we are now demanding that the model inputs themselves be the accurate outputs of processes at the genetic level that can be modeled by genomic prediction. It is not surprising, therefore, that modeling technologies (ranging from data collection to estimation) that were adequate for past applications now require improvement.

From a fundamental but traditional perspective, there are several issues of perennial concern in crop modeling. The first is model functional structure including both its degree of expressivity and its behavior under optimization. For example, estimation procedures like DE, that primarily yield point estimates, are limited in their ability to assess equifinality. At best, one can query the flatness of the goodness-of-fit function in the neighborhood of the estimate, but this does not tell anything about the ubiquity of equifinality across the parameter space. Nor do these procedures allow one to detect observations that fall outside of the model’s scope of expressivity unless the discrepancies are quite large. Doing so requires methods like the Sobol database scheme used here that can make broader assessments in both parameter and phenotype space. It may well be that the rarity with which database methods have been used has led to an underappreciation as to the prevalence of these adverse situations.

When expressivity issues are identified, results like those above are not likely to be solved merely by acquiring more data of the same type. In such situations, better models will often needed and modern genetic studies can help. A great many plant component subsystems are currently under study at the molecular level. Indeed, some of these (e.g., [61] are even being combined into multi-scale organ and whole plant models. Even without modeling directly at the genetic level one can use the derived insights to make informed choices between alternative representations of individual ecophysiological processes. Tardieu (2003) refers to such representations as “meta-mechanisms”. It would seem plausible that building models from component parts of increased biological realism should increase the ability to reproduce field variation - at the very least, it is hard to see how it can hurt. As a concrete example, the B73 parent is photoperiod insensitive. In CERES-Maize, however, the only way to express this is by setting P2O in excess of the observed photoperiods, with the consequences we have seen.

This is not to say, however, that both more and better data are not needed. Indeed, data quality issues can impact both expressivity and GSP stability. For example, while the date seed that are physically sown in a field is usually known and not subject to error, researchers often report a subjective notion of “effective sowing date” based on their interpretation of whether low soil moisture delayed germination. If errors in sowing date push an anthesis observation across the expressivity frontier, erroneous GSP estimates will result. Such errors can also arise if different personnel are involved across locations or growing seasons, especially for visually evaluated phenotypes like most phenological traits. Providing the emergence date can provide a partial check for these problems and also for errors in simulating time from sowing to emergence. Unfortunately, emergence dates were not reported for the maize NAM dataset.

Another traditional modeling concern has always been the relationship between the observed environmental data and the immediate environmental conditions actually experienced by individual plants. Weather data can suffer from multiple sources of bias and error [63]. For example, stations that are not located within or directly adjacent to experiments may have bias due to local variation in weather conditions. Additionally, although of limited concern for anthesis dates, the quality of soil and management data. In this study any systematic differences in protocols for collection of weather data between the sites as aggravated by small sample effects, might have contributed to some degree to the significance levels in Table 5. It would certainly be desirable to have a method by which this potential effect might be quantitatively assessed. Such a method could be instrumental in designing experimental procedures for reducing the problem. One potential example might be to eschew external measurements of some environmental variables (e.g., air temperature) and use sensors onboard UAV’s or other automated vehicles to measure plant temperatures or other critical features directly at high temporal and spatial frequencies.

More involved data types and structures are also needed to resolve issues of equifinality when they arise. Equifinality is fundamentally a problem of discernment. In simple terms, given an equation *c* = *a* + *b*, if one only has data on *c*, then estimates of *a* and *b* are doomed to be equifinal. If one desires otherwise, one must find a way to measure either *a* or *b*. Current technological efforts to develop high throughput phenotyping approaches might be quite helpful in this regard. For example, assuming that TOLN=P1/(PHINT×2)+5 is the correct way to model the number of leaves at anthesis, data on total leaf number would help constrain the parameter estimates. This leads toward a range of constrained and/or multiobjective estimation procedures on which there has been significant amounts of research [64,65]. Maximum entropy methods offer another opportunity wherein one identifies a probability distribution of values that is constrained by but mathematically no more informed than is justified by a set of potentially diverse data types [66]. Another alternative might be Bayesian methods with multivariate likelihood functions that combine several observational variables [67].

Another approach to reduce equifinality would be to use simpler models. The fewer the number of processes and GSP’s in a model, the smaller the opportunity for hard-to-spot tradeoffs to exist wherein adjustments to one parameter can be offset by tweaking another one. Of course, the tradeoff may be less expressivity leading to other problems. However, Welch et al. (2005) presented 12 dichotomies comparing gene network modeling and quantitative genetics approaches, where aspects of the former might also apply to ecophysiological modeling. They opined that an optimal modeling approach should entail a synthesis of both. The key features to be contributed from the network (i.e., ecophysiological) side would be (1) the ability to handle time-varying dynamics, (2) a far more parsimonious approach to expressing biological and biology × environmental interactions, and (3) a more mechanistic explanation of how traits originate. It is at least conceivable that some way station of moderate complexity exists between statistical genetics and full crop models that can achieve this.

At whatever level of complexity proves appropriate, one cannot accurately estimate the parameters controlling model components without collecting data on settings wherein the relevant processes operate differentially. This is clear from the P2O gap phenomenon, which was apparent when only short day data was used and absent under long days. Both settings distorted the results, in one case compressing estimates into a restricted range, leaving a gap, and, in the other, allowing them to spread out. Furthermore, this interacted with the range of values allowed, which caused shifts between (P1, PHINT) and (P2, P2O) as to which parameters appeared to be “explanatory”. The debilitating influence of such behavior on linking parameter values to genes is terribly obvious.

However, it also should not escape notice that the gap was evident even in a mixture of environments, suggesting that good experimental design entails more than just making sure that a suitable range of environments is included. There is some notion of balance that needs to be established and applied globally to data selection. In this context, it is worth noting that despite the fact that thousands of lines were planted in each location, there were only 539 lines where data were reported from all 11 trials. However, given the expense of such large-scale trials and the multiple purposes each one will serve, “balance” cannot mean “orthogonality” where all lines are planted at all sites. Of course, an established benefit of ecophysiological models is to serve as guides to help prioritize experimentation over time. It seems likely that as their integration with statistical genetic models expands, they might also be able to assist in the rational planning and resource allocation for large, multi-site trials.

Another approach entirely would be to seek to move beyond a two-step “estimate and then map” paradigm. Conventional mapping methods essentially isolate genetic markers whose pattern of assignment to lines mirrors the pattern of phenotype values of interest. A general linear model is assumed to mediate between marker states and realized phenotypes. There is no conceptual reason why that general linear model might not be replaceable by a crop model. In effect, one could conceive a hierarchical model in which a first-level model is specified on the data and higher order submodels are specified on the parameters that characterize the behavior of observed data, much like proposed by Bello et al. (2010).

One could conceptually implement this hierarchy in the context of crops by fitting phenotypes with an ECM whose GSP’s are then specified as functions of genetic markers at another level of the hierarchical model. Indeed, this is what the current paradigm attempts, except that the two-step estimation process curtails smooth borrowing of information across hierarchical levels of the model that could potentially help resolve the equifinality problem.

We acknowledge that one-step hierarchical model approach might not solve the sort of expressivity problems described in the thought experiment and documented in our results (both in section “Model expressivity”). Yet, it would enable the genetic structure of the population to inform the GSP estimation process. The potential utility of this hierarchical modeling approach is currently under study in one of our labs. The approach would also enable more efficient use of data. Currently, the two-step approach requires data from multiple environments [39] for each line in order to estimate the GSP’s before mapping can proceed. However, consider a line that was culled very early in the selection process, perhaps even after a single round. Because the parameters estimated in putative one-step hierarchical modeling schemes would include marker effects, even just one planting becomes a usable observation if the line is genotyped. This is a sufficiently inexpensive operation now that some programs (e.g. CIMMYT; [69] are doing so routinely for the offspring of all crosses.

A one-step hierarchical modeling approach might also make it possible to utilize data taken on lines after they enter the market place. Analogously to high throughput phenotyping in breeding programs, precision agricultural management is also investing in sensor-and model-based approaches to improve productivity [70,71] while collecting a wealth of multivariate data. Usually, of course, hybrids are released into areas where they show low G×E interactions. For example, a line with a particular P2O is not likely to be released across a sufficient range of latitudes to have great differences in day length. This would make it difficult to directly estimate P2O for the line using the methods described in this paper.

However, in a one-step hierarchical model approach, one would only be looking for markers that influenced P2O. In this case, data from many lines and geographical areas could be used together. This would also make such data usable for the sorts of hypothesis testing about genes discovered by other means, thus facilitating genetically-informed ecophysiological modeling. For such approaches to be workable, however, there are many policy issues to be resolved including information property rights and fair economic returns to data, not to mention the need to greatly harden cybersecurity protections [72]. However, if this can be done then issues of environmental coverage would likely be ameliorated due to the extent of the data that would become available.

## Conclusions

The original and seemingly simple goal of this study was to first fit the anthesis date component of the CERES-Maize model to data from over 5000 genotyped lines and then genetically map the resulting GSP values. However, we were unexpectedly detoured when we found that despite the high predictive quality of the values obtained, there were numerous artifacts that emerged in the estimation process, thereby making our immediate goal unachievable. We find it interesting that the problems we encountered would likely be invisible, though present, in smaller data sets and, unless addressed by suitable research, these problems bode ill for understanding any genetic underpinnings of ecophysiological models. This is worrisome given the recent escalating attention that has been given to this method of melding ecophysiological and statistical genetic models as a way of accelerating the crop improvement process so as to help meet global food and fiber needs by 2050.

The constraining issues fall into two categories. The first arises in situations where the model is unable to express the observed data for some line even by a relatively few days. In this circumstance, the line is assigned the GSP associated with the nearest point on model’s expression frontier – values which can, however, change only slowly along that boundary. The result is that many and in some cases a large majority of lines are assigned the same GSP values independent of their actual genetics.

The second symptom arises when the model can reproduce the data. In these instances, there can be many combinations of GSP values that predict equally well. When such equifinality exists, there is no principled way to assign the line a genetically relevant value. In short, when the model can express the data there is no unique combination of GSP values and, when unique combinations do exist they are often values being given to many lines because of a deficiency in model expressivity.

This finding is rather remarkable because in both breeding efforts and, indeed, genetic studies as a whole, anthesis date is considered, if not a simple trait, at least one that has proved much easier to elucidate than many others. In addition, it is generally, much more readily predicted by classical phenology models for reasons that, themselves, have become generally understood [37]. This cannot but make one wonder, what pitfalls might lie in wait for efforts to probe other, more involved traits.

Therefore, the next question to be asked by follow-on research is how prevalent are these phenomena. The best way to do that would seem to be to use Sobol database search methods. This is because, unlike optimizers that find single “best estimates”, the database approach will reveal the both the extent of the expressible phenotype regions as well as a direct measure of the extent of any equifinality.

However, despite the ability to reuse results databases for many searches, undertaking such a program in any broadly based fashion will be highly demanding computationally. For this reason, strong consideration should be given to disaggregating comprehensive models into separate modules that can be studied independently at much lower computational cost. (This is what we did for the limited DE run, although Python certainly is not a high performance language.) A better long-term strategy would be to program future models in a manner that supports single-module testing at the source code level. Doing so will facilitate the whole-model verifications needed to ensure that fragmentation into modules for testing and improvement by different labs does not compromise integration at the level of the scientific community.

As module testing and innovation progress, it will be of strategic value to ground improvements in advancing genetic understanding at the molecular level. While this might seem daunting to those versed in purely physiological approaches, it need not be so. One of the most venerable concepts in all of the life sciences is that of the biological hierarchy that is, a series of many functional levels extending from molecules to the biosphere. One of the perspectives emerging from molecular science is that that hierarchy might, be operationally much flatter than commonly believed. That is, simple changes at lower levels can easily create tangible responses multiple levels higher. To the extent that this is true, it greatly reduces the complexity of bridging across those levels. This is the philosophy behind the meta-mechanism approach mentioned earlier [62,73].

That approach has a proven ability to account for environmental interactions with sufficient skill to eliminate observed G×E interactions from GSP’s in the data sets used (Reymond et al., 2003). However, as shown by the *p*-values in Table 5, the very large data set used herein conveyed an extraordinary power to detect site-year dependencies in GSP estimation. Indeed, so powerful as to make one wonder if an insignificant result is scientifically achievable by any even remotely feasible research effort? A better number to use for practical evaluations might be the index of variability in Table 5. This would give a clear index of the size of the effect as a percentage of the parameter values. Also, means exist for comparing such indices to see if reductions in their values (i.e. by an improved model with lowered site-year set dependency) are statistically significant (Vangel, 1996).

A final message from our research is that one cannot fix problems that one does not know exist. Community interest in the fitting-and-mapping paradigm has been high as shown by the heavy citation rates for the seminal papers in this area. For example, as of September, 2016, the Hammer et al. (2006) paper had been cited 257 times and those publications, *themselves*, had been cited by 6,370 others (Source: Google Scholar). There is also no doubt as to the importance of the ability to predict the behaviors of novel genotypes in novel environments while crosses are still in the planning stage. Indeed, this is precisely the genotype-to-phenotype problem, which has been declared by the National Research Council to be a top-priority goal for applied biology (NRC, 2008). So these impediments need to be overcome. However, with methods now in hand to detect adverse model behaviors under estimation, research that is probing ever more deeply into the control mechanisms of plant growth and development, and concrete tests to document model improvements, there is no reason to believe that we cannot do so.

## Acknowledgements

The plan to use the maize NAM data to first developed through discussions with iPlant (http://www.iplantcollaborative.org) on novel applications of cyberinfrastructure in plant science. The authors acknowledge the Texas Advanced Computing Center (TACC; http://www.tacc.utexas.edu) at The University of Texas at Austin and Beocat, Kansas State University for providing high performance computing resources that have contributed to the research results reported within this paper. Support for this effort was also supplied by the Department of Agronomy at Kansas State University.

## Author Contributions

Conceptualization: JW KT SW. Methodology: AL SW KT JW. Analyzed the data: AL SW. Manuscript preparation: AL SW JW KT

